# Exploitation of ATP-sensitive potassium ion (KATP) channels by HPV promotes cervical cancer cell proliferation by contributing to MAPK/AP-1 signalling

**DOI:** 10.1101/2022.09.22.508991

**Authors:** James A. Scarth, Christopher W. Wasson, Molly R. Patterson, Debra Evans, Diego Barba-Moreno, Holli Carden, Adrian Whitehouse, Jamel Mankouri, Adel Samson, Ethan L. Morgan, Andrew Macdonald

## Abstract

Persistent infection with high-risk human papillomaviruses (HPVs) is the causal factor in multiple human malignancies, including >99% of cervical cancers and a growing proportion of oropharyngeal cancers. Prolonged expression of the viral oncoproteins E6 and E7 is necessary for transformation to occur. Although some of the mechanisms by which these oncoproteins contribute to carcinogenesis are well-characterised, a comprehensive understanding of the signalling pathways manipulated by HPV is lacking. Here, we present the first evidence to our knowledge that the targeting of a host ion channel by HPV can contribute to cervical carcinogenesis. Through the use of pharmacological activators and inhibitors of ATP-sensitive potassium ion (K_ATP_) channels, we demonstrate that these channels are active in HPV-positive cells and that this activity is required for HPV oncoprotein expression. Further, expression of SUR1, which forms the regulatory subunit of the multimeric channel complex, was found to be upregulated in both HPV+ cervical cancer cells and in samples from patients with cervical disease, in a manner dependent on the E7 oncoprotein. Importantly, knockdown of SUR1 expression or K_ATP_ channel inhibition significantly impeded cell proliferation via induction of a G1 cell cycle phase arrest. This was confirmed both *in vitro* and in *in vivo* tumourigenicity assays. Mechanistically, we propose that the pro-proliferative effect of K_ATP_ channels is mediated via the activation of a MAPK/AP-1 signalling axis. A complete characterisation of the role of K_ATP_ channels in HPV-associated cancer is now warranted in order to determine whether the licensed and clinically available inhibitors of these channels could constitute a potential novel therapy in the treatment of HPV-driven cervical cancer.

## Introduction

It has been estimated that high-risk human papillomaviruses (HPVs) are the causal factor in over 5% of all human cancers, including >99.7% of cervical cancers and a growing number of oropharyngeal cancers [1, 2]. HPV16 is responsible for the majority of these (around 55% of cervical cancers and almost all HPV-positive (HPV+) head and neck cancers), whilst HPV18 is the cause of another 15% of cervical cancers [3]. As a result of this, cervical cancer is the fourth most prevalent cancer in women and the most common cause of cancer-related death in young women [1].

HPV-associated malignancies are the result of a persistent infection where the host immune system fails to detect and clear the virus efficiently, although even in this situation carcinogenesis may take several years to occur [1]. The most important factor required for initiation and progression of cancer is the prolonged increased expression of the viral oncoproteins E6 and E7, which results in the dysregulation of cell proliferation [4]. Some of the mechanisms by which the oncoproteins achieve this have been widely-studied, including the ability of HPV E7 to drive S phase re-entry via binding to and inducing the degradation of pRb and the related pocket proteins p107 and p130 [5–7]. Concurrently, E6 targets p53 for proteasomal degradation by recruiting the E3 ubiquitin ligase E6-associated protein (E6AP) in order to block pro-apoptotic signalling [8]. Further, high-risk E6 proteins are also able to increase telomerase activity and bind to and regulate PSD95/DLG/ZO-1 (PDZ) domain-containing proteins in order to increase cell proliferation and survival [9, 10], whilst E7 has a key role in mediating evasion of the host immune response [11, 12]. More recently, E6 has been shown to modulate a multitude of host signalling pathways, including the JAK-STAT and Hippo pathways, during transformation [13–18].

However, a comprehensive understanding of the host signalling networks modulated by HPV during transformation is still lacking. Furthermore, no therapeutic targeting of HPV-associated proteins in HPV-driven malignancies currently exists. Therefore, it is necessary to identify novel HPV-host interactions and to establish whether they may constitute potential new therapeutic targets. In particular, despite the availability of prophylactic vaccines, there are currently no effective anti-viral drugs for use against HPV. Current therapeutics rely on the widely used yet non-specific DNA-damaging agent cisplatin in combination with radiotherapy [19, 20]. However, resistance to cisplatin, either intrinsic or acquired, is a significant problem [21]. Although this issue can be somewhat alleviated through the use of combination therapy involving cisplatin alongside paclitaxel, there is an urgent need to develop more targeted therapies for the treatment of HPV-associated malignancies [22].

Ion channels may represent ideal candidates for these novel therapies given the abundance of licensed and clinically available drugs targeting the complexes which could be repurposed if demonstrated to be effective [23]. Indeed, the importance of ion channels in the regulation of the cell cycle and cell proliferation has become increasingly recognised [24–28]. Cells undergo a rapid hyperpolarisation during progression through the G1-S phase checkpoint, which is then reversed during G2 [26]. It is thought that potassium ion (K^+^) channels are particularly important for this, with a number of K^+^ efflux channels having been observed to be increased in expression and activity during G1 [25, 26]. Of these, ATP-sensitive K^+^ (K_ATP_) channels have been shown to be expressed highly in some cancers, and channel inhibition can result in decreased proliferation [29–33].

K_ATP_ channels are hetero-octameric membrane complexes consisting of four pore-forming Kir6.x subunits (either Kir6.1 or Kir6.2) surrounded by four regulatory sulfonylurea receptor (SUR) subunits [34]. Multiple isoforms of the sulfonylurea receptor exist: SUR1 (encoded by *ABCC8*), and SUR2A and SUR2B, which are produced via alternative splicing of the *ABCC9* transcript [35]. K_ATP_ channels are expressed in multiple tissues, although the composition of the channels can vary, which may account for subtle tissue-specific properties of the channels [34].

In this study, we performed a pharmacological screen to identify potassium ion channels that may play a role in HPV pathogenesis. We identified that, of the K^+^ channels investigated, inhibition of K_ATP_ channels had a negative impact on HPV oncoprotein expression and cellular transformation. By screening for the expression of K_ATP_ channel subunits, we identified that the SUR1 subunit is expressed highly in HPV+ cervical cancer cells, and that this increased SUR1 expression is driven by the E7 oncoprotein. Depletion of K_ATP_ channel activity, either by siRNA-mediated knockdown or pharmacological inhibition, significantly impeded proliferation and cell cycle progression. Further, we propose that this pro-proliferative effect is mediated via the activation of a mitogen-activated protein kinase (MAPK)/activator protein-1 (AP-1) signalling axis. We hope that the targeting of K_ATP_ channels may prove to be beneficial in the treatment of HPV-associated neoplasia.

## Results

### K_ATP_ channels are important for HPV gene expression in cervical cancer cells and primary human keratinocytes

Ion channels are emerging as crucial regulators of cell signalling pathways and cell cycle progression [24–27]. In particular, K^+^ channels have been shown to be active or expressed highly in a variety of cancer cell lines [36, 37]. Furthermore, a growing number of viruses have been shown to be capable of modulating the activity of host ion channels [38]. To determine whether HPV requires the activity of K^+^ channels either during its life cycle or during transformation, we performed a pharmacological screen using several broadly acting K^+^ channel modulators. We first assessed the impact of K^+^ channel inhibition on the expression of the viral oncoprotein E7, which is essential for the survival and proliferation of cervical cancer cells both *in vitro* and *in vivo* [39–41]. Treatment of either HPV18+ cervical cancer cells (HeLa) or primary human keratinocytes containing the HPV18 genome with tetraethylammonium (TEA), quinine or quinidine at pharmacologically relevant concentrations resulted in a decrease in E7 oncoprotein expression (**Fig 1A**), indicating that HPV does indeed require the activity of K^+^ channels. Addition of potassium chloride (KCl) to increase the extracellular K^+^ concentration and thus collapse the plasma membrane potential had the same effect on oncoprotein expression, but addition of sodium chloride (NaCl) had no impact, indicating that alteration of osmolarity alone does not impact upon HPV gene expression (**Fig 1A**).

**Fig 1.**
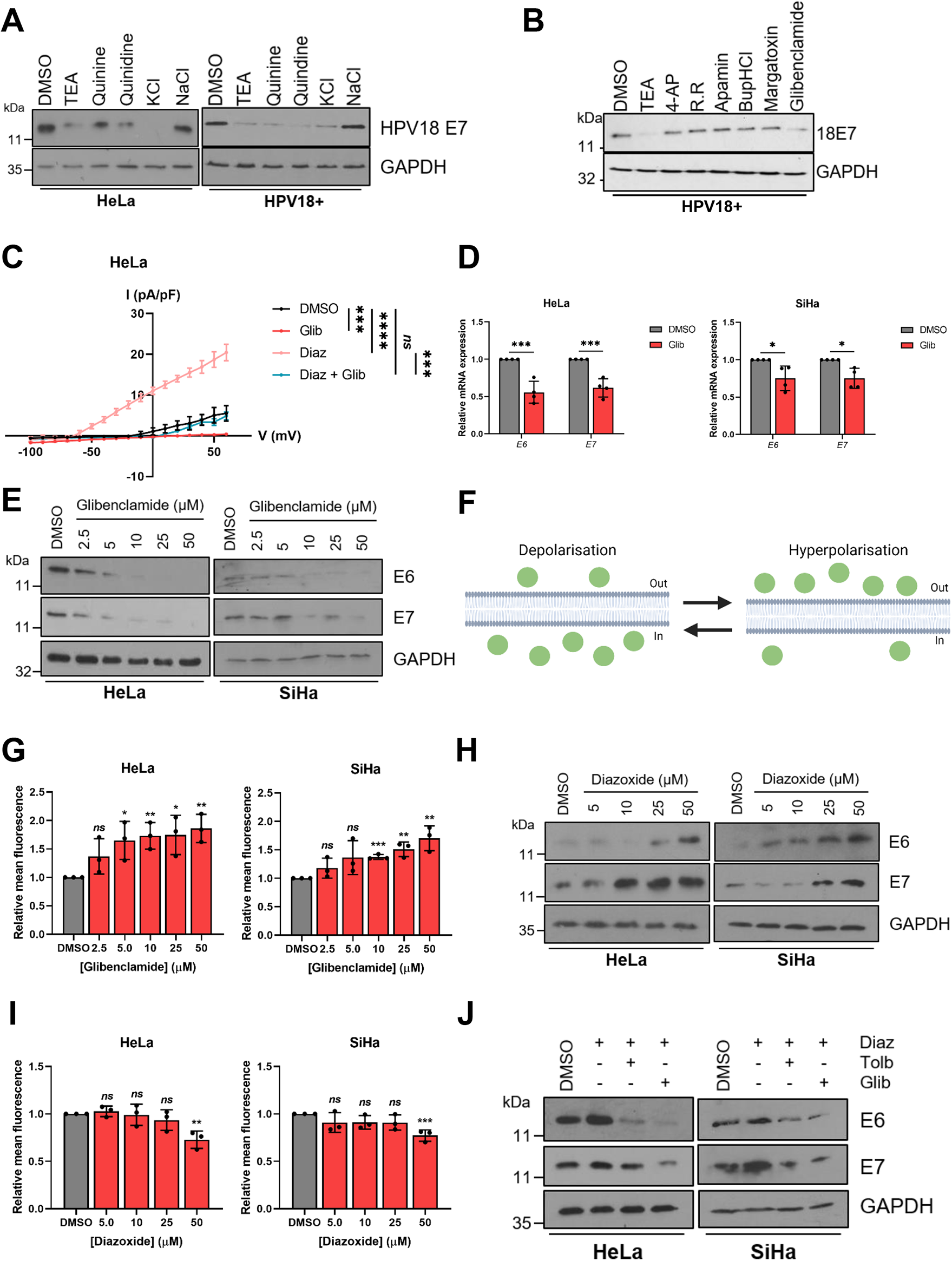
K_ATP_ channels are important for HPV gene expression. **A)** Representative western blots for E7 expression in HeLa cells and primary human keratinocytes containing HPV18 episomes treated with DMSO, a broadly acting K^+^ channel inhibitor (25 mM tetraethylammonium (TEA), 100 µM quinine, 100 µM quinidine or 70 mM KCl) or 70 mM NaCl. GAPDH served as a loading control. **B)** Representative western blots for E7 expression in HPV18+ primary keratinocytes treated with DMSO, 25 mM TEA or one of a panel of class-specific K^+^ channel inhibitors (2 mM 4-aminopyridine (4-AP), 50 µM ruthenium red (RR), 10 nM apamin, 20 µM bupivacaine hydrochloride (BupHCl), 10 nM margatoxin or 50 μM glibenclamide). GAPDH served as a loading control. **C)** Mean current density-voltage relationships for K^+^ currents in HeLa cells treated with DMSO, diazoxide (50 μM), glibenclamide (10 μM), or both diazoxide and glibenclamide (n = 5 for all treatments). **D)** Expression levels of *E6* and *E7* mRNA in HeLa and SiHa cells treated with glibenclamide (10 μM) measured by RT-qPCR. Samples were normalised against *U6* mRNA levels. **E)** Representative western blots of E6 and E7 expression in HeLa and SiHa cells treated with increasing doses of glibenclamide. GAPDH served as a loading control. **F)** Schematic illustrating the plasma membrane permeability of DiBAC_4_(3). Figure created using BioRENDER.com. **G)** Mean DiBAC_4_(3) fluorescence levels in HeLa and SiHa cells treated with increasing dose of glibenclamide. Samples were normalised to DMSO controls. **H)** Representative western blots of E6 and E7 expression in HeLa and SiHa cells serum starved for 24h (to reduce basal E6/E7 expression) prior to treatment with increasing doses of diazoxide. GAPDH served as a loading control. **I)** Mean DiBAC_4_(3) fluorescence levels in HeLa and SiHa cells treated with increasing dose of diazoxide. Samples were normalised to DMSO control. **J)** Representative western blots of E6 and E7 expression in HeLa and SiHa cells treated with diazoxide (50 μM) alone or in combination with glibenclamide (10 μM) or tolbutamide (200 μM). Bars represent means ± standard deviation (SD) of a minimum of three biological replicates with individual data points displayed where possible. *Ns* not significant, *P<0.05, **P<0.01, ***P<0.001 (Student’s t-test).

In order to identify whether a particular class of K^+^ channels was required for HPV gene expression, a further screen was performed using more specific K^+^ channel inhibitors. Treatment with inhibitors of voltage-gated K^+^ channels (4-aminopyridine (4-AP) or margatoxin), two-pore-domain K^+^ channels (ruthenium red (RR) or bupivacaine hydrochloride (BupHCl)) or calcium-activated K^+^ channels (apamin) had no impact on E7 protein levels (**Fig 1B**). However, treatment with glibenclamide, a clinically available inhibitor of K_ATP_ channels which acts via binding to the regulatory SUR subunits [42], resulted in a marked decrease in E7 expression in HPV18 containing primary keratinocytes (**Fig 1B**).

Next, electrophysiological analysis was performed in HeLa cells to confirm that active K_ATP_ channels were present. A clear outward K^+^ current was observed, which was significantly increased upon application of the K_ATP_ channel activator diazoxide (**Fig 1C**). This was entirely reversed after addition of the channel inhibitor glibenclamide, whilst glibenclamide treatment alone was able to reduce basal K^+^ currents. Together, this confirms that active K_ATP_ channels are present in cervical cancer cells.

To further investigate the repressive effects of glibenclamide on the HPV oncoproteins, expression levels were assayed after treatment of both HPV16+ and HPV18+ cervical cancer cells with the inhibitor at a range of concentrations. A significant decrease in expression of both E6 and E7 was observed at both the mRNA level (**Fig 1D**) and the protein level (**Fig 1E**) at concentrations as low as 10 μM. To ensure that the effect of glibenclamide treatment on viral oncoprotein expression was due to the inhibition of K_ATP_ channel activity, we first analysed the membrane potential of cells using the fluorescent dye Bis-(1,3-Dibutylbarbituric Acid) Trimethine Oxonol (DiBAC_4_(3)) [43, 44]. The ability of the dye to enter cells is proportional to the degree to which the plasma membrane is depolarised (**Fig 1F**). Therefore, the dose-dependent increase in fluorescence observed following glibenclamide treatment indicates an increasing level of depolarisation, consistent with a reduction in K_ATP_ channel opening (**Fig 1G**). Significantly, treatment of cells with tolbutamide, a member of the same class of sulfonylurea drugs as glibenclamide, also resulted in a dose-dependent decrease in oncoprotein expression with a corresponding increase in DiBAC_4_(3) fluorescence (**S1A-C Fig**).

In line with our inhibitor data, treatment of HPV+ cervical cancer cells with the K_ATP_ channel activator diazoxide resulted in a dose-dependent increase in HPV oncoprotein expression (**Fig 1H**). As before, analysis of DiBAC_4_(3) fluorescence was performed to assess the impact of diazoxide treatment on the plasma membrane potential. A decrease in fluorescence, particularly apparent at the highest concentration of 50 µM, was observed after application of diazoxide, indicating increasing levels of hyperpolarisation (**Fig 1I**). Finally, treatment of HPV+ cervical cancer cells with either glibenclamide or tolbutamide abolished the diazoxide-induced increase in HPV oncoprotein expression (**Fig 1J**). Taken together, these data demonstrate that K_ATP_ channel activity is important in the regulation of HPV gene expression.

### HPV upregulates the SUR1 subunit of K_ATP_ channels

Given the importance of K_ATP_ channel activity for HPV oncoprotein expression, we hypothesised that HPV may upregulate expression of channel subunits. Notably, the expression of K_ATP_ channel subunits displays significant tissue-specific variability, so it was important to gain an understanding of which isoforms are expressed in cervical tissue [45]. We therefore screened for the expression of all K_ATP_ channel subunits in a panel of cervical cancer cell lines by RT-qPCR. K_ATP_ channels are hetero-octameric complexes consisting of four pore-forming Kir6.x subunits (either Kir6.1 or Kir6.2) surrounded by four regulatory SURx subunits (either SUR1, SUR2A or SUR2B) [34]. We found no significant difference in the expression of Kir6.1 (*KCNJ8*), whilst Kir6.2 (*KCNJ11*) expression was higher in HPV-negative (HPV-) C33A cells, as well as three of the four HPV+ cervical cancer cell lines, when compared with normal human keratinocytes (NHKs) (**Fig 2A**). Expression of SUR2A (*ABCC9A*) could not be detected in any of the cell lines, and SUR2B (*ABCC9B*) was not significantly increased in any of the HPV+ cell lines relative to NHKs. However, expression of the SUR1 (*ABCC8*) subunit was significantly higher in all four of the HPV+ cancer cell lines examined, with no increase detected in HPV-C33A cells. We therefore focussed on the SUR1 subunit for the purposes of this study.

**Fig 2.**
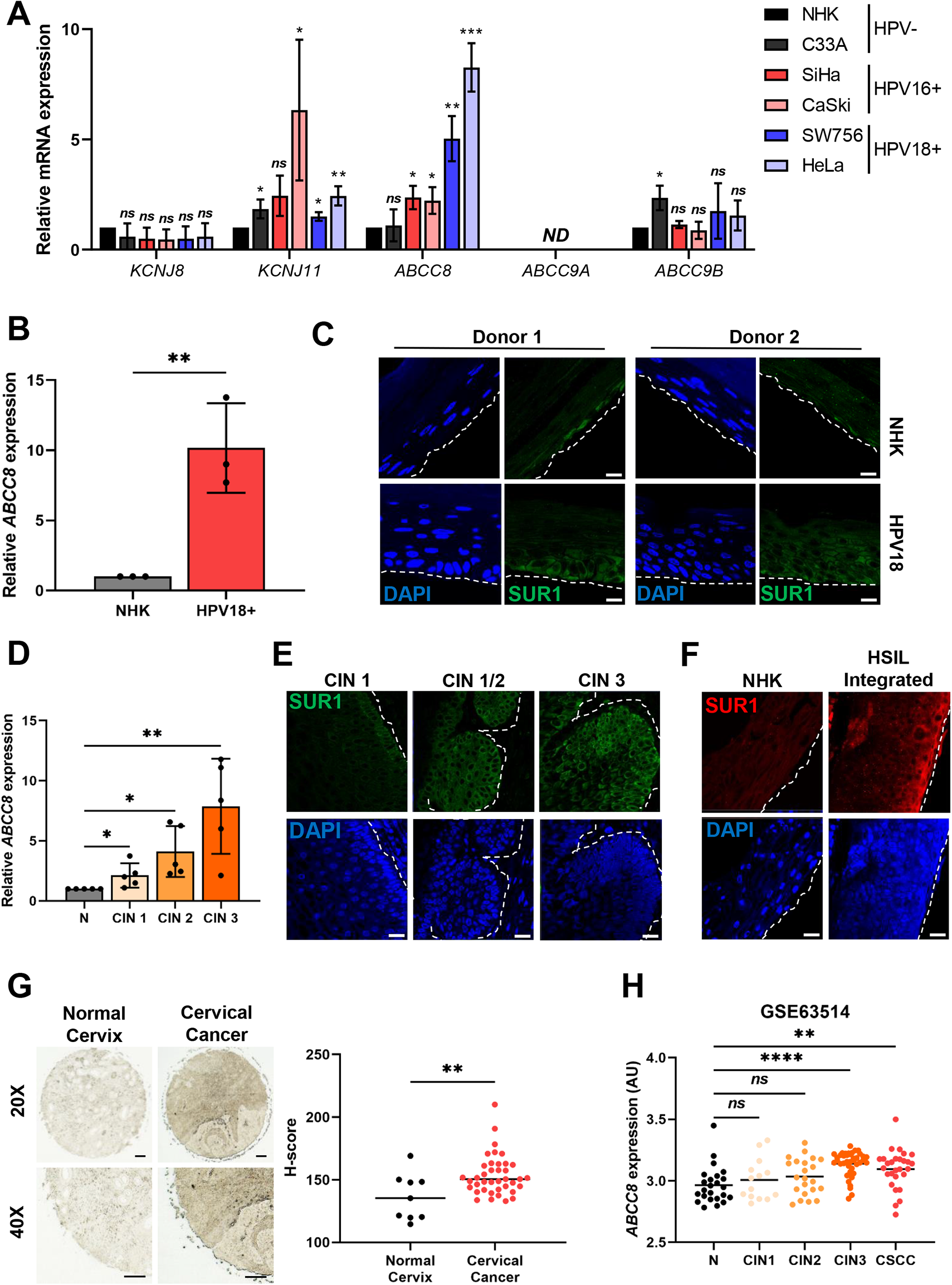
HPV enhances expression of the SUR1 subunit of K_ATP_ channels. **A)** mRNA expression levels of K_ATP_ channel subunits in HPV-normal human keratinocytes (NHK) and a panel of five cervical cancer cell lines – one HPV- (C33A), two HPV16+ (SiHa and CaSki), and two HPV18+ (SW756 and HeLa) detected by RT-qPCR. Samples were normalised against *U6* mRNA levels. Data is displayed relative to NHK controls. **B)** *ABCC8* mRNA expression in NHKs and keratinocytes containing episomal HPV18 genomes detected by RT-qPCR. Samples were normalised against *U6* mRNA levels. **C)** Representative immunofluorescence analysis of sections from organotypic raft cultures of NHK and HPV18+ keratinocytes detecting SUR1 levels. Nuclei were visualised with DAPI and the white dotted line indicates the basal layer. Two donor cell lines were used to exclude donor-specific effects. Images were acquired with identical exposure times. Scale bar, 40 μm. **D)** *ABCC8* mRNA expression in a panel of cervical cytology samples representing normal cervical tissue (N) and cervical disease of increasing severity (CIN 1 - 3) detected by RT-qPCR (n = 5 from each grade). Samples were normalised against *U6* mRNA levels. **E)** Representative immunofluorescence analysis of tissue sections from cervical lesions of increasing CIN grade. Sections were stained for SUR1 levels (green) and nuclei were visualised with DAPI (blue). Images were acquired with identical exposure times and the white dotted line indicates the basal layer. Scale bar, 40 μm. **F)** Representative immunofluorescence analysis of sections from organotypic raft cultures of NHK and a W12 cell line presenting with HSIL morphology detecting SUR1 levels (red). Nuclei were visualised with DAPI (blue) and the white dotted line indicates the basal layer. Images were acquired with identical exposure times. Scale bar, 40 μm. **G)** Representative immunohistochemistry analysis and scatter dot plots of quantification of normal cervical (n = 9) and cervical cancer (n = 39) tissue sections stained for SUR1 protein. Scale bar, 50 μm. **H)** Scatter dot plot of expression data acquired from the GSE63514 dataset. Arbitrary values for *ABCC8* mRNA expression in normal cervix (n = 24), CIN1 lesions (n = 14), CIN2 lesions (n = 22), CIN3 lesions (n = 40) and cervical cancer (n = 28) samples were plotted. Bars represent means ± SD of a minimum of three biological replicates with individual data points displayed where possible. *Ns* not significant, *P<0.05, **P<0.01, ***P<0.001, ****P<0.0001 (Student’s t-test).

To further investigate the increased SUR1 expression potentially induced by HPV, we analysed cell lines containing episomal HPV18 generated from primary foreskin keratinocytes. Transcript levels of *ABCC8* (SUR1) were significantly increased by approximately 10 fold relative to the NHK control (**Fig 2B**). Additionally, sections of organotypic raft cultures of NHKs and HPV18-containing keratinocytes, which recapitulate all stages of the HPV life cycle [46], were analysed for SUR1 protein levels by immunofluorescence microscopy. This demonstrated a marked increase in SUR1 protein expression in the suprabasal layers of HPV18+ rafts in comparison to NHK raft cultures, consistent across both donors (**Fig 2C**). Next, SUR1 expression was analysed in cervical liquid-based cytology samples from a cohort of HPV16+ patients representing the progression of cervical disease development (CIN 1 – CIN 3). We observed an increase in *ABCC8* (SUR1) mRNA levels relative to HPV-normal cervical tissue which correlated with disease progression, with the highest expression observed in CIN 3 samples (**Fig 2D**). Indeed, immunofluorescence microscopy analysis of human cervical sections classified as LSIL (CIN 1), LSIL with foci of HSIL (CIN 1/2) and HSIL (CIN 3), confirmed that SUR1 protein levels increase with cervical disease progression (**Fig 2E**).

Furthermore, SUR1 protein levels were analysed using the HPV16+ W12 *in vitro* model system [13, 47]. At low passage numbers, these cells display an LSIL phenotype in raft culture, but long-term passaging results in a phenotype more closely mirroring that of HSIL and squamous cell carcinoma. Raft cultures were generated from NHKs and a W12 clone representing a HSIL phenotype and stained for SUR1 protein levels. High levels of SUR1 staining were observed in the HSIL raft compared to the NHK control (**Fig 2F**), thus confirming that SUR1 expression increases with cervical disease progression. In order to analyse SUR1 protein levels in cervical cancer tissue, we performed immunohistochemistry using an array of normal cervix and cervical cancer tissue sections. Significantly higher SUR1 expression was observed in the cancer tissue sections, as indicated by an increase in H-score (**Fig 2G**). Finally, to confirm our above observations, we mined an available microarray database containing data from primary cervical disease and tumour samples. This revealed a significant increase in *ABCC8* (SUR1) expression in the CIN 3 and cervical squamous cell carcinoma (CSCC) samples (**Fig 2H**). Taken together, these data indicate that SUR1 expression is increased in HPV-containing keratinocytes and, importantly, in HPV-associated cervical disease.

### Depletion of SUR1 reduces HPV gene expression in cervical cancer cells

After identifying that the SUR1 subunit of K_ATP_ channels was highly expressed during HPV+ cervical disease, we investigated the effects of suppressing SUR1 expression. Knockdown of SUR1 using a pool of specific siRNAs was performed in both HPV16+ (SiHa) and HPV18+ (HeLa) cervical cancer cells (**Fig 3A**). Furthermore, monoclonal HeLa cell lines stably expressing one of two SUR1-specific shRNAs were also generated (**Fig 3B**). To ascertain the effect of SUR1 depletion on the plasma membrane potential, DiBAC_4_(3) fluorescence was used. We observed a ~2 fold increase in fluorescence after siRNA treatment, indicating a significant depolarisation characteristic of a reduction in K_ATP_ channel activity (**Fig 3C**). We were unable to analyse the impact of stable suppression of SUR1 on the plasma membrane potential due to the presence of a ZsGreen selectable marker.

**Fig 3.**
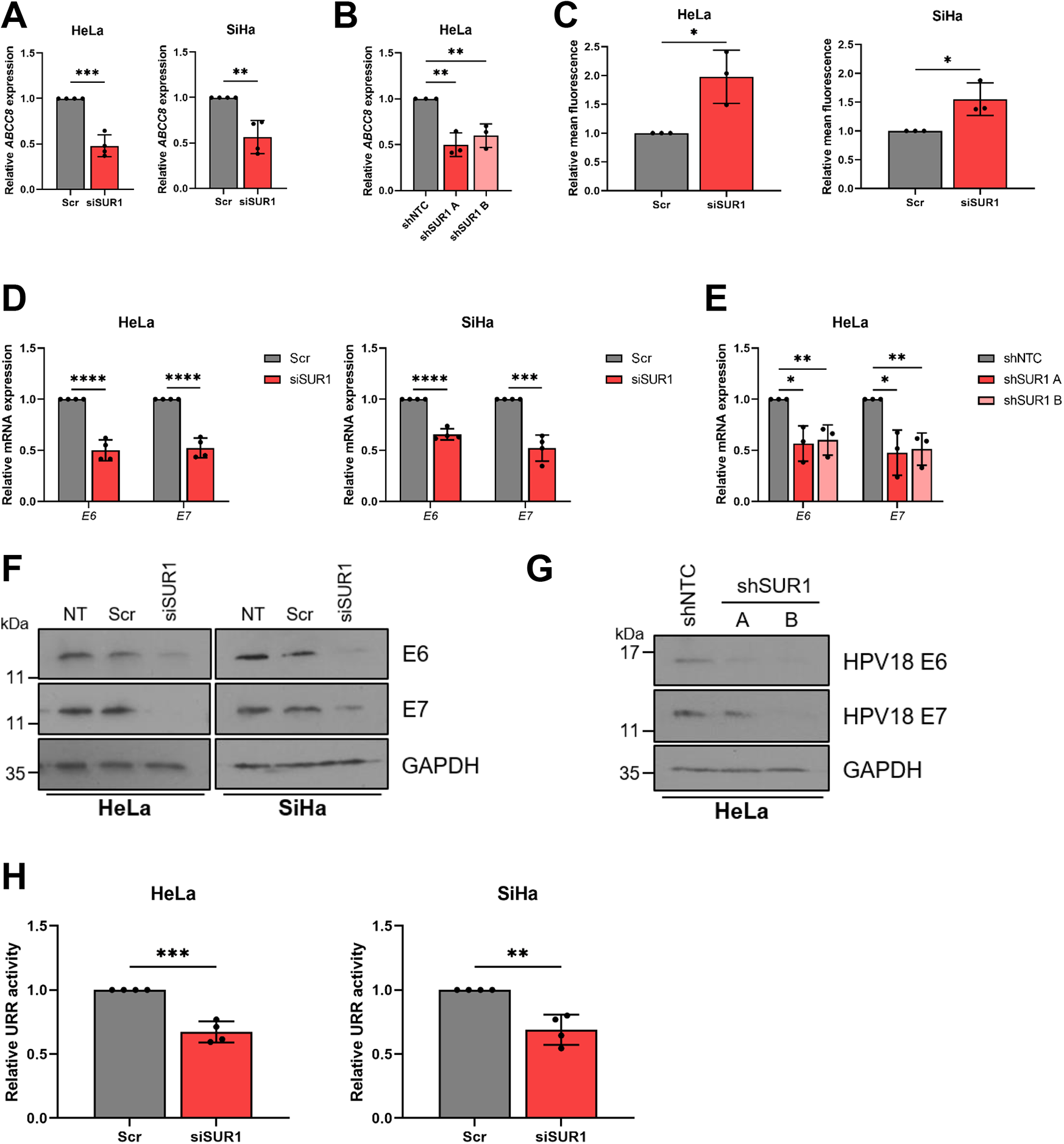
Depletion of SUR1 reduces HPV gene expression in cervical cancer cells. **A-B)** Relative expression of *ABCC8* mRNA in **A)** HeLa and SiHa cells transfected with a pool of SUR1-specific siRNA and **B)** two monoclonal HeLa cell lines stably expressing SUR1-specific shRNAs measured by RT-qPCR. Samples were normalised against *U6* mRNA levels. **C)** Relative mean DiBAC_4_(3) fluorescence levels in HeLa and SiHa cells transfected with SUR1 siRNA. Samples were normalised to scramble controls. **D-E)** Relative expression of *E6* and *E7* mRNA in **D)** HeLa and SiHa cells transfected with SUR1 siRNA and **E)** HeLa cell lines stably expressing SUR1-specific shRNAs measured by RT-qPCR. Samples were normalised against *U6* mRNA levels. **F-G)** Representative western blots of E6 and E7 expression in **F)** HeLa and SiHa cells transfected with SUR1 siRNA and **G)** HeLa cell lines stably expressing either a non-targeting (shNTC) or a SUR1-specific shRNA. GAPDH served as a loading control. **H)** Relative firefly luminescence in HeLa and SiHa cells co-transfected with SUR1 siRNA and either a HPV18 or HPV16 URR reporter plasmid respectively. Luminescence values were normalised against *Renilla* luciferase activity and data is displayed relative to scramble controls. Bar graphs represent means ± SD of a minimum of three biological replicates with individual data points displayed. *P<0.05, **P<0.01, ***P<0.001, ****P<0.0001 (Student’s t-test).

Subsequently, the effect of SUR1 depletion on HPV gene expression was analysed. siRNA-mediated knockdown of SUR1 resulted in a significant decrease in HPV oncoprotein expression, measured both at the transcript and protein level (**Fig 3D and 3F**). The same impact on HPV gene expression was also observed following stable knockdown of SUR1, when compared to cells expressing a non-targeting shRNA (shNTC) (**Fig 3E and 3G**). To confirm that the effect of SUR1 depletion on HPV gene expression was due to a direct loss of transcription from the viral upstream regulatory region (URR), luciferase reporters containing the HPV16 and HPV18 URRs were used. We observed a significant decrease in relative luciferase activity after SUR1 knockdown with both URR reporter plasmids, confirming a direct loss of HPV early promoter activity (**Fig 3H**).

In contrast to this, transfection of a pool of SUR2-specific siRNAs had no impact on HPV oncoprotein expression in either HPV16+ or HPV18+ cervical cancer cells, in line with our data showing that HPV does not induce an increase in expression of the SUR2 subunit of K_ATP_ channels (**S2A-2D Fig**). We did however observe a small depolarisation of the plasma membrane, indicated by an increase in DiBAC_4_(3) fluorescence (**S2B Fig**), suggesting that a small minority of K_ATP_ channels in HPV+ cervical cancer cells may be composed of the SUR2 subunit.

Finally, in order to confirm that the effects on HPV gene expression observed were K_ATP_ channel-dependent, rather than a potential channel-independent function of SUR1, Kir6.2 levels were also depleted using a pool of specific siRNAs (**S3A Fig**). The ~2 fold and ~1.5 fold increases in DiBAC_4_(3) fluorescence resulting from Kir6.2 knockdown observed in HeLa and SiHa cells respectively were broadly in line with the changes observed following SUR1 knockdown (**S3B Fig**). Knockdown of Kir6.2 resulted in a significant reduction in both mRNA and protein levels of E6 and E7 to a similar extent to that detected following SUR1 knockdown (**S3C and 3D Fig**), thus confirming that the impacts of SUR1 depletion were indeed channel-dependent. Taken together, these data confirm that K_ATP_ channel expression in cervical cancer cells is important for HPV oncoprotein expression.

### The E7 oncoprotein is responsible for the increased SUR1 expression in HPV+ cervical cancer cells

We next explored the mechanism behind the observed HPV-induced increases in SUR1 expression. Given that the E6 and E7 oncoproteins are key drivers of transformation, we hypothesised that they may be responsible for the heightened SUR1 levels [2]. To investigate this, expression of both the E6 and E7 oncoproteins was repressed using siRNA in HPV+ cervical cancer cell lines. We saw a >70% decrease in *ABCC8* (SUR1) mRNA levels after knockdown of oncoprotein expression (**Fig 4A**). A decrease in *ABCC8* mRNA expression could also be observed after silencing of E6 and E7 in HPV18+ primary keratinocytes (**Fig 4B**), indicating that oncoprotein expression is necessary to induce SUR1 expression. In order to gain an understanding of which oncoprotein drives this, the E6 and E7 oncoproteins of HPV18 were overexpressed in turn and in combination in HPV-C33A cells. HPV18 E6 did not result in any change in *ABCC8* mRNA levels, whereas in contrast, expression of E7 led to a ~2.5 fold increase in *ABCC8* expression (**Fig 4C**). Co-expression of E6 alongside E7 did not cause a further increase in *ABCC8* mRNA levels, indicating that the E7 oncoprotein is the major driver of SUR1 expression. Similar effects on *ABCC8* mRNA levels were observed when this was performed in NHKs (**Fig 4D**). Further, C33A cell lines stably expressing HA-tagged HPV18 oncoproteins were generated as previously described [18]. A significant upregulation of *ABCC8* expression was only observed in the HA-E7 expressing cells, consistent with our transient overexpression data (**Fig 4E**).

**Fig 4.**
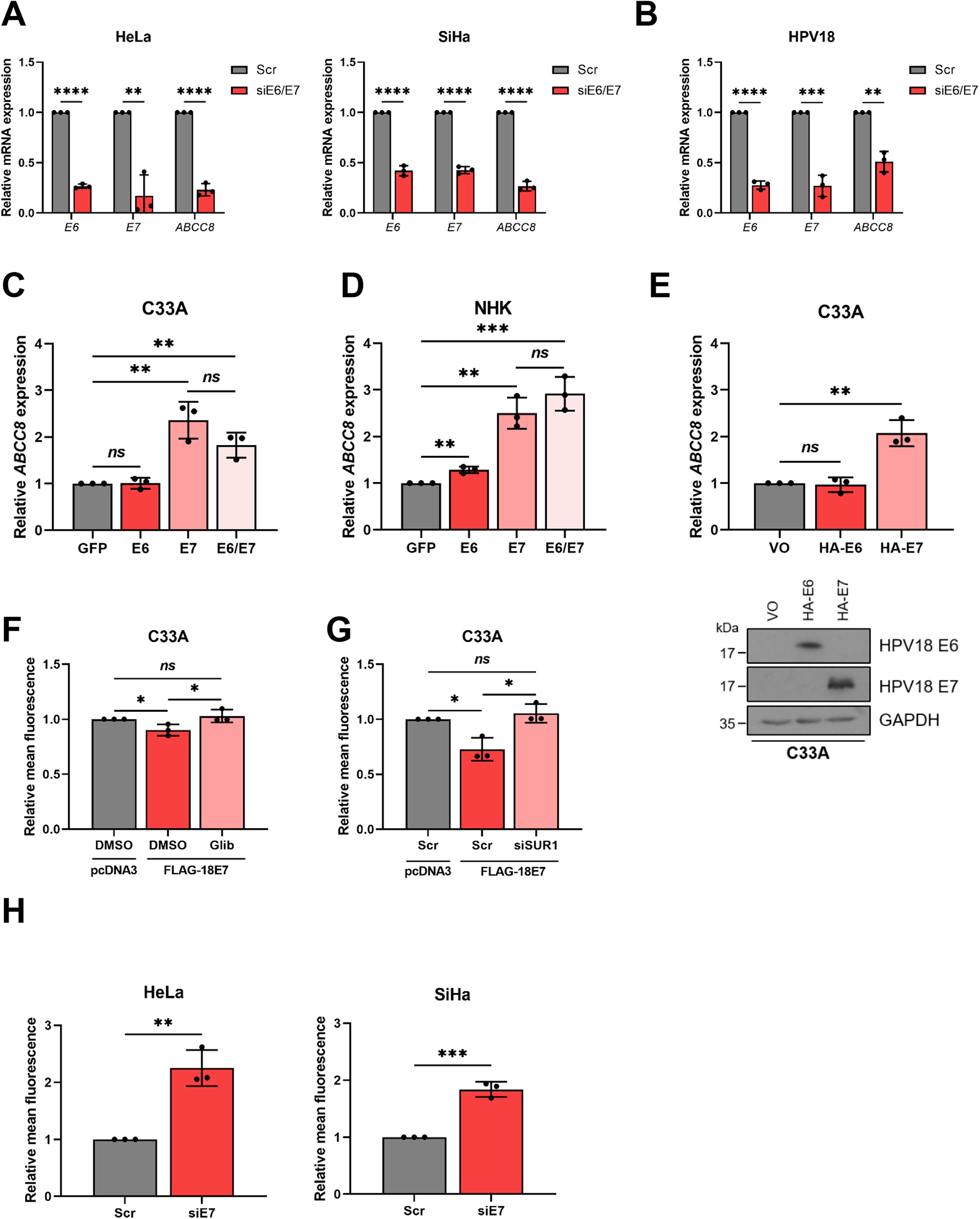
The E7 oncoprotein is responsible for the increase in SUR1 expression. **A-B)** Relative *ABCC8* mRNA expression measured by RT-qPCR in **(A)** HeLa and SiHa cells and **(B)** HPV18+ primary keratinocytes co-transfected with E6- and E7-specific siRNA. Samples were normalised against *U6* mRNA levels. Successful knockdown was confirmed by analysing *E6* and *E7* mRNA levels. **C-D)** Expression levels of *ABCC8* mRNA measured by RT-qPCR in **(C)** C33A cells and **(D)** NHKs transfected with GFP-tagged HPV18 oncoproteins. Samples were normalised against *U6* mRNA levels and data is presented relative to the GFP-transfected control. Successful transfection was confirmed by immunofluorescence analysis (not shown). **E)** Relative expression of *ABCC8* mRNA in C33A cells stably-expressing HA-tagged HPV18 oncoproteins measured by RT-qPCR. Samples were normalised against *U6* mRNA levels. Expression of oncoproteins was confirmed by western blot. **F)** Mean DiBAC_4_(3) fluorescence levels in C33A cells after transfection of FLAG-tagged HPV18 E7 and treatment with either DMSO or glibenclamide (10 μM). Samples were normalised to the pcDNA3-transfected control. **G)** Mean DiBAC_4_(3) fluorescence levels in C33A cells after co-transfection of FLAG-tagged HPV18 E7 and SUR1-specific siRNA. Samples were normalised to the pcDNA3/scramble-transfected control. **H)** Mean DiBAC_4_(3) fluorescence levels in HeLa and SiHa cells after transfection of HPV E7-specific siRNA. Samples were normalised to the scramble control. Bars represent means ± SD of a minimum of three biological replicates with individual data points displayed. *Ns* not significant, *P<0.05, **P<0.01, ***P<0.001, ****P<0.0001 (Student’s t-test).

To confirm that the observed changes in SUR1 expression led to an impact on K_ATP_ channel activity, the plasma membrane potential of cells was assayed after overexpression of HPV18 E7 in HPV-cervical cancer cells. A significant reduction in DiBAC_4_(3) fluorescence, indicative of membrane hyperpolarisation, was detected (**Fig 4F and 4G**). This decrease was abolished both by treatment with glibenclamide and by siRNA-mediated knockdown of SUR1, suggesting that the hyperpolarisation was due to an E7-dependent increase in K_ATP_ channel activity. Additionally, silencing of HPV E7 expression resulted in a ~2 fold increase in DiBAC_4_(3) fluorescence (**Fig 4H**), consistent with a reduction in K_ATP_ channel opening. These data indicate that the E7 oncoprotein, rather than E6, is the major factor regulating HPV-induced increases in SUR1 expression.

### K_ATP_ channels drive proliferation in HPV+ cervical cancer cells

Given the effects of K_ATP_ channel activity on HPV gene expression, it was hypothesised that modulation of channel activity may too impact upon the proliferation of HPV+ cervical cancer cells. Treatment of both HPV16+ SiHa cells and HPV18+ HeLa cells with glibenclamide to inhibit K_ATP_ channel activity resulted in a significant decrease in cell proliferation (**Fig 5A**), anchorage-dependent (**Fig 5B**) and anchorage-independent colony formation (**Fig 5C**). In contrast, treatment of HPV-C33A cells with glibenclamide had minimal impact on proliferation or colony-forming ability (**Fig 5A-C**).

**Fig 5.**
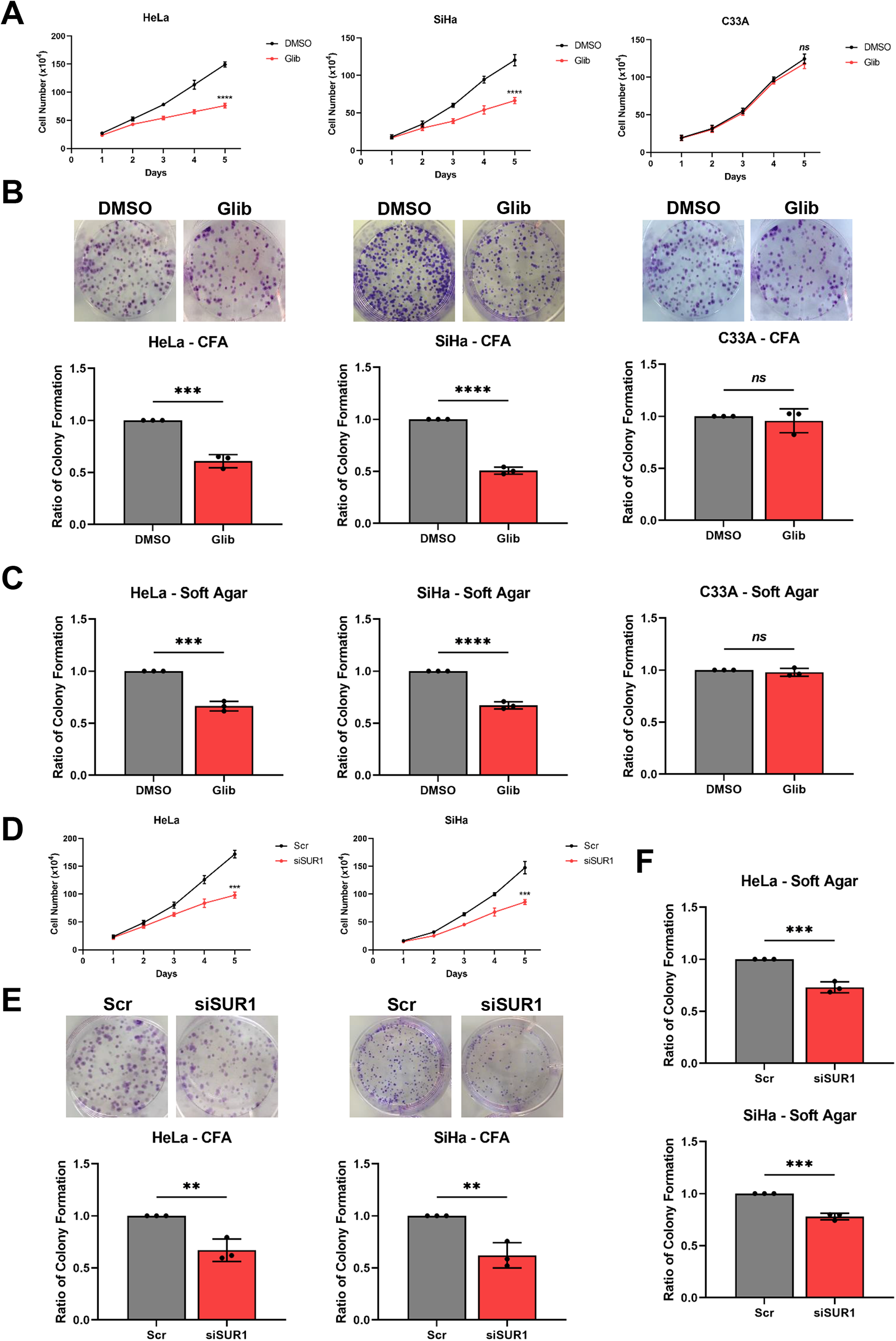
K_ATP_ channels drive proliferation in cervical cancer cells. **A-C)** Growth curve analysis **(A)**, colony formation assay (to measure anchorage-dependent growth) **(B)** and soft agar assay (to measure anchorage-independent growth) **(C)** of HeLa, SiHa and C33A cells after treatment with DMSO or glibenclamide (10 μM) for 24 hours. **D-F)** Growth curve analysis **(D)**, colony formation assay **(E)** and soft agar assay **(F)** of HeLa and SiHa cells after transfection of SUR1-specific siRNA. Data shown is means ± SD of a minimum of three biological replicates with individual data points displayed where appropriate. *Ns* not significant, *P<0.05, **P<0.01, ***P<0.001, ****P<0.0001 (Student’s t-test).

To eliminate the possibility of off-target effects of glibenclamide, a pool of SUR1-specific siRNAs was used to confirm the impact of reduced K_ATP_ channel activity on HPV+ cervical cancer cell proliferation. Both HPV16+ and HPV18+ cells demonstrated a reduced proliferation rate (**Fig 5D**) and colony-forming ability (**Fig 5E and 5F**) after depletion of SUR1 levels. Further, stable suppression of SUR1 levels via the expression of specific shRNAs resulted in a similar or greater impact on proliferation, and the anchorage-dependent and anchorage-independent colony-forming ability of HPV18+ cervical cancer cells (**S4A-C Fig**). In contrast, depletion of the alternative K_ATP_ channel regulatory subunit SUR2 had a minimal impact on the proliferation and colony-forming ability of both HPV+ cervical cancer cell lines analysed (**S2E-G Fig**). This is consistent with our data showing a lack of an effect on HPV oncoprotein expression after silencing of SUR2.

Finally, to confirm that the reduction in proliferation observed after either glibenclamide treatment or suppression of SUR1 expression was a result of decreased K_ATP_ channel activity, we analysed the growth of HPV+ cervical cancer cells following siRNA depletion of the pore-forming Kir6.2 subunit. This resulted in a decrease in proliferation and colony formation in both HPV16+ and HPV18+ cervical cancer cells, concordant with that observed following SUR1 depletion (**S3E-G Fig**). Collectively, these data demonstrate that K_ATP_ channels are important drivers of proliferation in HPV+ cervical cancer cells.

### K_ATP_ channel overexpression is sufficient to stimulate proliferation in the absence of HPV

Given the emerging evidence indicating that K_ATP_ channel activation can promote proliferation, and that heightened channel expression has been demonstrated in some other cancer types [29–33], we hypothesised that overexpression of K_ATP_ channel subunits alone (i.e. in the absence of HPV) may be sufficient to increase proliferation of cervical cancer cells. The individual subunits were therefore overexpressed alone and in combination in HPV-C33A cervical cancer cells. Expression of Kir6.2 alone had no impact on the proliferation or colony forming ability of the cells (**Fig 6A-C**). Whilst SUR1 overexpression did result in a small increase in anchorage-dependent and anchorage-independent colony formation, a significantly greater increase was observed when both subunits were overexpressed in combination (**Fig 6A-C**). Together, these data indicate that K_ATP_ channel activity is pro-proliferative, and that the reduction in cell growth observed herein following channel inhibition or knockdown is likely not solely due to a loss of HPV oncoprotein expression.

**Fig 6.**
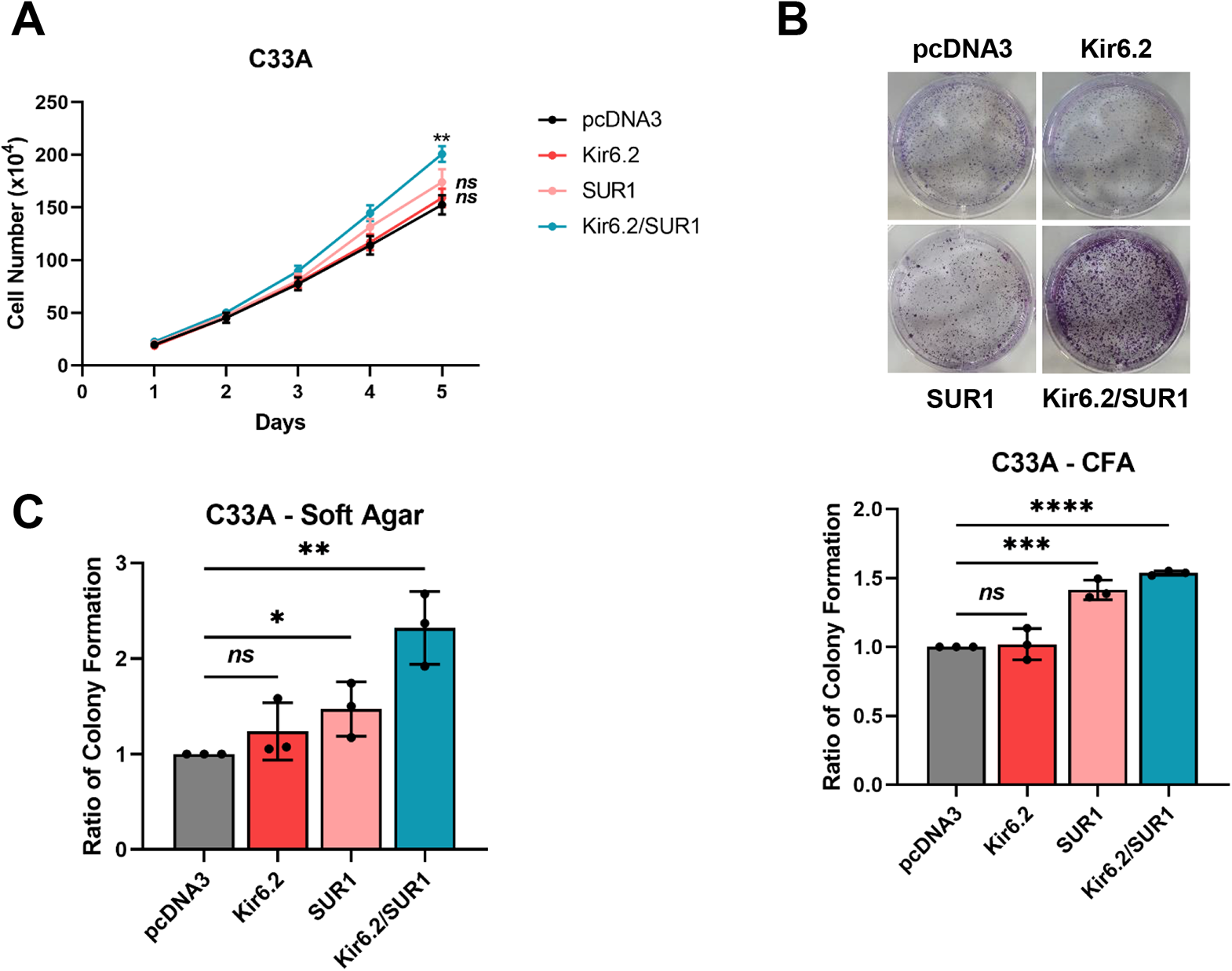
K_ATP_ channel overexpression is sufficient to stimulate proliferation in the absence of HPV. Growth curve analysis **(A)**, colony formation assay **(B)** and soft agar assay **(C)** of C33A cells transfected with plasmids expressing HA-tagged Kir6.2 and/or SUR1. Data shown is means ± SD of three biological replicates with individual data points displayed where appropriate. *Ns* not significant, *P<0.05, **P<0.01, ***P<0.001, ****P<0.0001 (Student’s t-test).

### K_ATP_ channel activity regulates progression through the G1/S phase transition

To further evaluate the impact of reduced K_ATP_ channel activity on cell proliferation, we assessed the cell cycle distribution of HPV+ cervical cancer cells using flow cytometry after blockade of K_ATP_ channel activity. We felt this to be particularly pertinent as K_ATP_ channel inhibition has been shown to result in a G1 phase arrest in glioma and breast cancer cell lines [30, 31]. In line with this, a significant increase in the proportion of cells in G1 phase was observed after both pharmacological inhibition of channel activity and siRNA-mediated SUR1 silencing in both HPV16+ and HPV18+ cervical cancer cells (**Fig 7A and 7B**). As cyclins are key regulators of cell cycle progression, we also analysed their expression after glibenclamide treatment or suppression of SUR1 levels. We detected significant decreases in expression at both the mRNA and protein level of cyclin D1 (*CCND1*) and cyclin E1 (*CCNE1*), both of which regulate progression through G1 and the transition into S phase (**Fig 7C-F**). This was consistent across both cell lines and treatments. In contrast, we observed only minimal changes in cyclin A2 (*CCNA2*) levels and no effect on cyclin B1 (*CCNB1*) expression at the mRNA level (**Fig 7C and 7D**), with similar observations at the protein level (**Fig 7E and 7F**). Together, these data suggest that K_ATP_ channels drive proliferation by regulating the G1/S phase transition.

**Fig 7.**
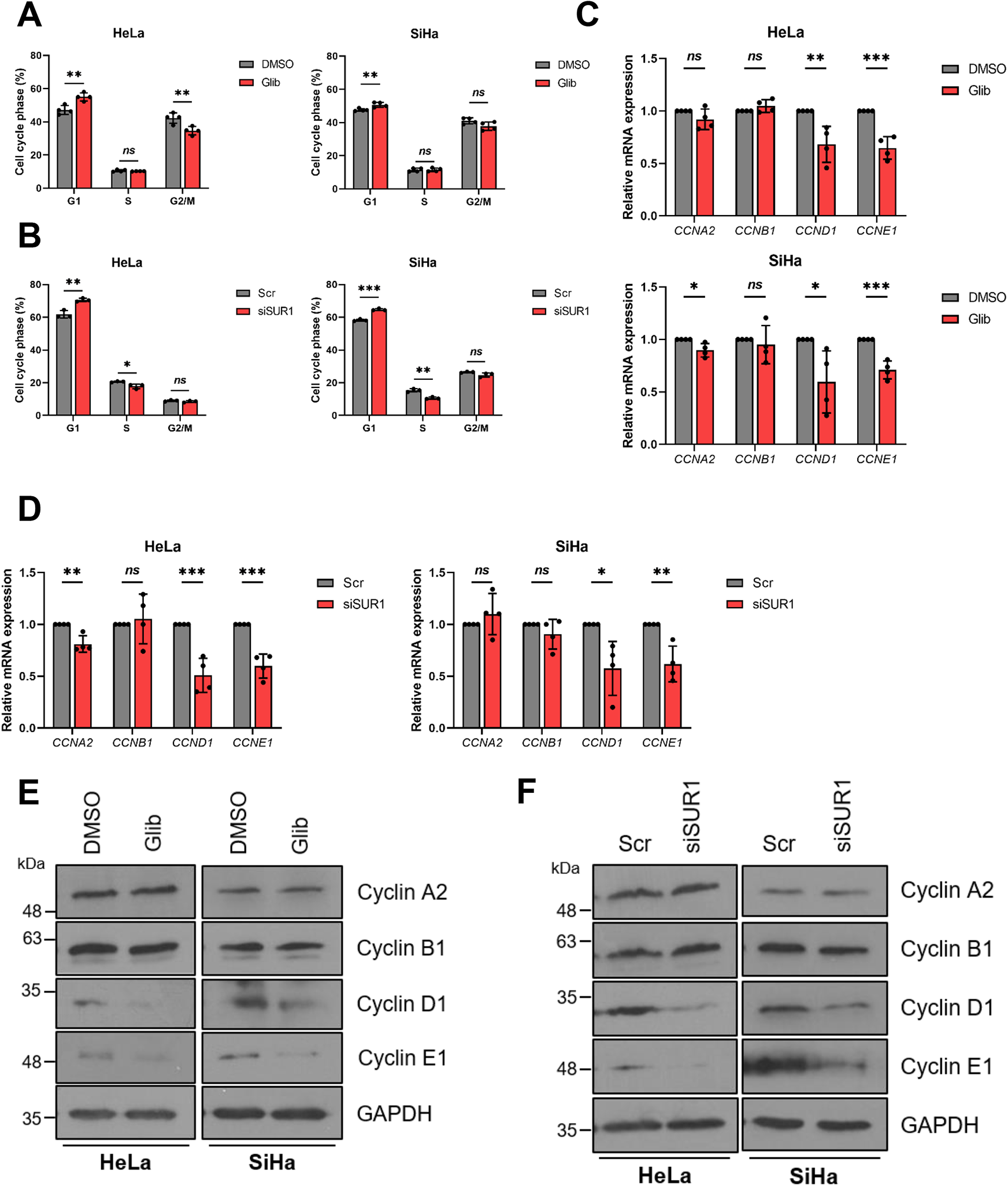
K_ATP_ channel activity regulates progression through the G1/S phase transition. **A-B)** Flow cytometry analysis of cell cycle phase distribution of HeLa and SiHa cells after **(A)** treatment with either DMSO or glibenclamide (25 μM) for 48 hours and **(B)** transfection of SUR1-specific siRNA. **C-D)** mRNA expression levels of cyclins in HeLa and SiHa cells after **C)** treatment with either DMSO or glibenclamide (10 μM) for 24 hours or **D)** transfection of SUR1-specific siRNA measured by RT-qPCR. Samples were normalised against *U6* mRNA levels and data is displayed relative to the appropriate control. **E-F)** Representative western blots of cyclin proteins in HeLa and SiHa cells after **E)** treatment with either DMSO or glibenclamide (10 μM) for 24 hours or **F)** transfection of SUR1-specific siRNA. GAPDH served as a loading control. Bars represent means ± SD of a minimum of three biological replicates with individual data points displayed. *Ns* not significant, *P<0.05, **P<0.01, ***P<0.001 (Student’s t-test).

### K_ATP_ channels are not required for the survival of cervical cancer cells

We also wanted to investigate the impact of channel inhibition on the survival of HPV+ cervical cancer cells. Increased levels of apoptosis have been reported in some cancer types after K_ATP_ channel blockade [29, 31, 32]. However, we failed to detect any increase in the cleavage of either poly(ADP) ribose polymerase (PARP) or caspase 3, both key indicators of the induction of apoptosis (**S5A Fig**). Further, we also failed to observe an increase in either early or late apoptosis via Annexin V staining of exposed phosphatidylserine on the plasma membrane (**S5B Fig**), thus confirming that K_ATP_ channel inhibition alone does not impact upon the survival of HPV+ cervical cancer cells.

### K_ATP_ channels contribute towards the activation of MAPK/AP-1 signalling

We next wanted to gain an understanding into the mechanism by which K_ATP_ channels promote proliferation in HPV+ cervical cancer cells. MAPK signalling is known to be a crucial driver of cell proliferation [48], and K_ATP_ channel opening can lead to activation of the MAP kinase ERK1/2 [49, 50], so we therefore analysed ERK1/2 phosphorylation levels following stimulation of HPV+ cervical cells with diazoxide. This revealed a significant increase in ERK1/2 phosphorylation post-stimulation, which was reversed following the addition of the MEK1/2 inhibitor U0126 (**Fig 8A**). Additionally, an increase in HPV18 E7 protein levels was observed, consistent with prior experiments; this was also reduced with U0126 treatment. Interestingly, an increase in both the phosphorylation and total protein expression of the AP-1 family member cJun was observed. AP-1 transcription factors are composed of dimers of proteins belonging to the Jun, Fos, Maf and ATF sub-families, and can regulate a wide variety of cellular processes, including proliferation, survival and differentiation [51]. This indicates that cJun/AP-1 could be a downstream target of ERK1/2 following K_ATP_ channel stimulation.

**Fig 8.**
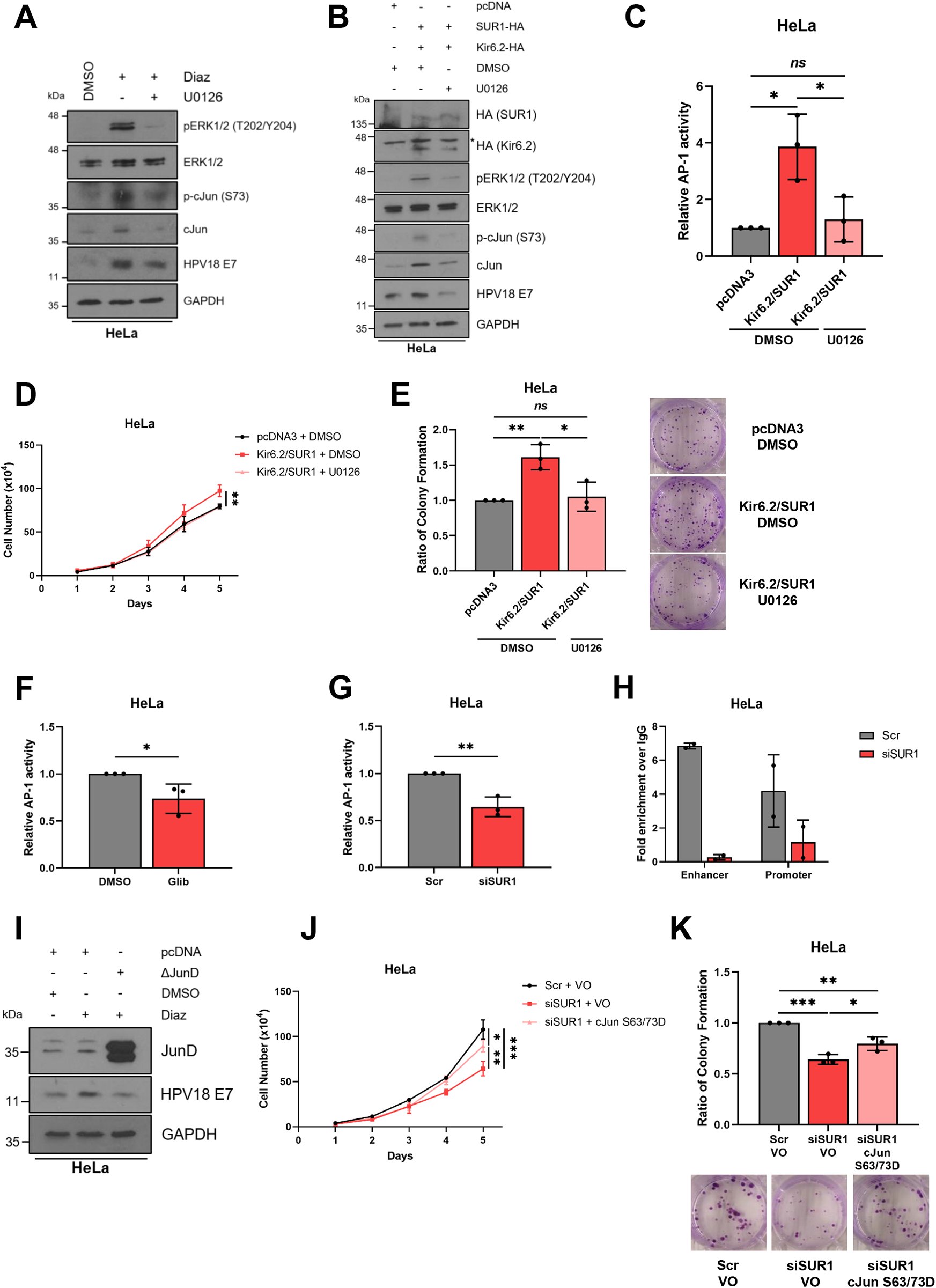
K_ATP_ channels drive proliferation by contributing towards the activation of MAPK/AP-1 signalling. **A-B)** Representative western blots of phospho-ERK1/2, ERK1/2, phospho-cJun, cJun and E7 in HeLa cells either **A)** serum starved for 24 hours prior to treatment with diazoxide (50 μM), with and without the MEK1/2 inhibitor U0126 (20 μM), for 24 hours or **B)** transfected with plasmids expressing HA-tagged Kir6.2 and SUR1, with and without U0126 treatment (20 μM). * denotes the presence of a non-specific band. GAPDH served as a loading control. **C)** Relative firefly luminescence in HeLa cells co-transfected with plasmids expressing HA-tagged Kir6.2 and SUR1 and an AP-1-driven reporter construct. Cells were also treated with DMSO or the MEK1/2 inhibitor U0126 (20 μM) for 24 hours. Luminescence values were normalised against *Renilla* luciferase activity. **D-E)** Growth curve analysis **(D)** and colony formation assay **(E)** of HeLa cells after co-transfection with plasmids expressing HA-tagged Kir6.2 and SUR1 and treatment with DMSO or U0126 (20 μM) for 24 hours. **F-G)** Relative firefly luminescence in HeLa cells transfected with an AP-1-driven reporter plasmid and either **F)** treated with glibenclamide (10 μM) or **G)** transfected with SUR1-specific siRNA. Luminescence values were normalised against *Renilla* luciferase activity and data is displayed relative to the appropriate control. **H)** ChIP-qPCR analysis of cJun binding to the HPV18 URR in HeLa cells transfected with SUR1-specific siRNA. Chromatin was prepared from HeLa cells and cJun immunoprecipitated using an anti-cJun antibody, followed by qPCR using primers specific to AP-1 binding sites in the HPV18 URR. cJun binding is presented as a fold increase over IgG binding (n = 2). **I)** Representative western blots for JunD, E6 and E7 expression in HeLa cells treated with diazoxide (50 μM), with and without transfection of a plasmid expressing ΔJunD. Cells were serum-starved for 24 hours prior to treatment. GAPDH served as a loading control. **J-K)** Growth curve analysis **(J)** and colony formation assay **(K)** of HeLa cells after co-transfection with SUR1-specific siRNA and a plasmid expressing a constitutively-active form of cJun (S63/73D). Bars represent means ± SD of a minimum of three biological replicates (unless stated otherwise) with individual data points displayed where appropriate. *Ns* not significant, *P<0.05, **P<0.01, ***P<0.001 (Student’s t-test).

To confirm these observations, overexpression of both K_ATP_ channel subunits in combination was performed. This similarly resulted in increased ERK1/2 phosphorylation, increases in both cJun phosphorylation and total protein levels, as well as enhanced E7 expression (**Fig 8B**). As before, these increases were reversed, in part, by the addition of U0126. To confirm that the changes in expression and phosphorylation of cJun corresponded to alterations in AP-1 activity, we employed a luciferase reporter construct containing three tandem AP-1 binding sites [52, 53]. K_ATP_ channel overexpression led to a ~4 fold increase in relative AP-1 activity, which was significantly reduced in the presence of U0126 (**Fig 8C**).

Following this, we performed assays to answer the question of whether the pro-proliferative effects of K_ATP_ channels are mediated by this MAPK/AP-1 signalling axis. We observed an increase in the proliferation and colony-forming ability of HeLa cells following overexpression of K_ATP_ channel subunits, which was reversed through MEK1/2 inhibition (**Fig 8D and 8E**).

Next, we investigated the impact of reducing K_ATP_ channel activity on AP-1 activity. Concordant with earlier data, a ~30% reduction in AP-1 activity was observed in HeLa cells following either glibenclamide treatment or transfection of SUR1-specific siRNA, as measured using a luciferase reporter assay (**Fig 8F and 8G**). To investigate whether modulation of K_ATP_ channel activity affected recruitment of cJun/AP-1 to the HPV URR, we performed ChIP-qPCR analysis using primers spanning the two AP-1 binding sites within the HPV18 URR, one in the enhancer region and one in the promoter [53]. This revealed that SUR1 knockdown reduced cJun recruitment to both binding sites within the viral URR, highlighting the critical role K_ATP_ channels may have in regulating oncoprotein expression (**Fig 8H**).

To further confirm our observations, we employed a dominant-negative JunD construct (ΔJunD): this encodes a truncated form of JunD which is able to dimerise with other AP-1 family members, yet lacks a transcriptional activation domain. Previous studies in our lab have validated that ΔJunD expression almost completely abolishes AP-1 activity [53]. Transfection of this construct resulted in a decrease in diazoxide-induced HPV oncoprotein expression (**Fig 8I**).

Finally, we examined whether the reintroduction of active cJun would be able to rescue the proliferation defect of HeLa cells transfected with SUR1-specific siRNA. This revealed a significant increase in both proliferation and colony-forming ability following expression of a constitutively-active form of cJun, in which the two key phosphorylatable residues S63 and S73 are mutated to aspartic acid to mimic phosphorylation (S63/73D) (**Fig 8J and 8K**) [54–56]. The rescue was incomplete however, illustrating that cJun/AP-1 is likely one of multiple targets downstream of K_ATP_ channel-induced ERK1/2 signalling. Taken together, these data indicate that K_ATP_ channel activity activates MAPK and AP-1 signalling to drive proliferation and oncoprotein expression.

### K_ATP_ channels drive proliferation *in vivo*

To confirm our *in vitro* observations, we performed tumourigenicity experiments using SCID mice. Animals were subcutaneously injected with HeLa cells stably expressing either a non-targetting shRNA or a SUR1-specific shRNA. Tumour development was monitored, revealing rapid growth in the HeLa shNTC control group, as expected (**Fig 9A**). However, a significant delay in the growth of tumours in all mice injected with SUR1 knockdown cells compared to the shNTC controls was observed (**Fig 9A**). To quantify this delay in growth, the period of time between injection of tumours and growth to a set volume (250 mm^3^) was calculated. This revealed that the SUR1-depleted tumours took an additional 11 days on average to reach an equivalent size (**Fig 9B**). Further, animals bearing SUR1-depleted tumours displayed significantly prolonged survival, with one mouse remaining alive at the conclusion of the study (**Fig 9C**). Together, these data demonstrate that K_ATP_ channels drive the growth of HPV+ cervical cancer cell xenografts.

**Fig 9.**
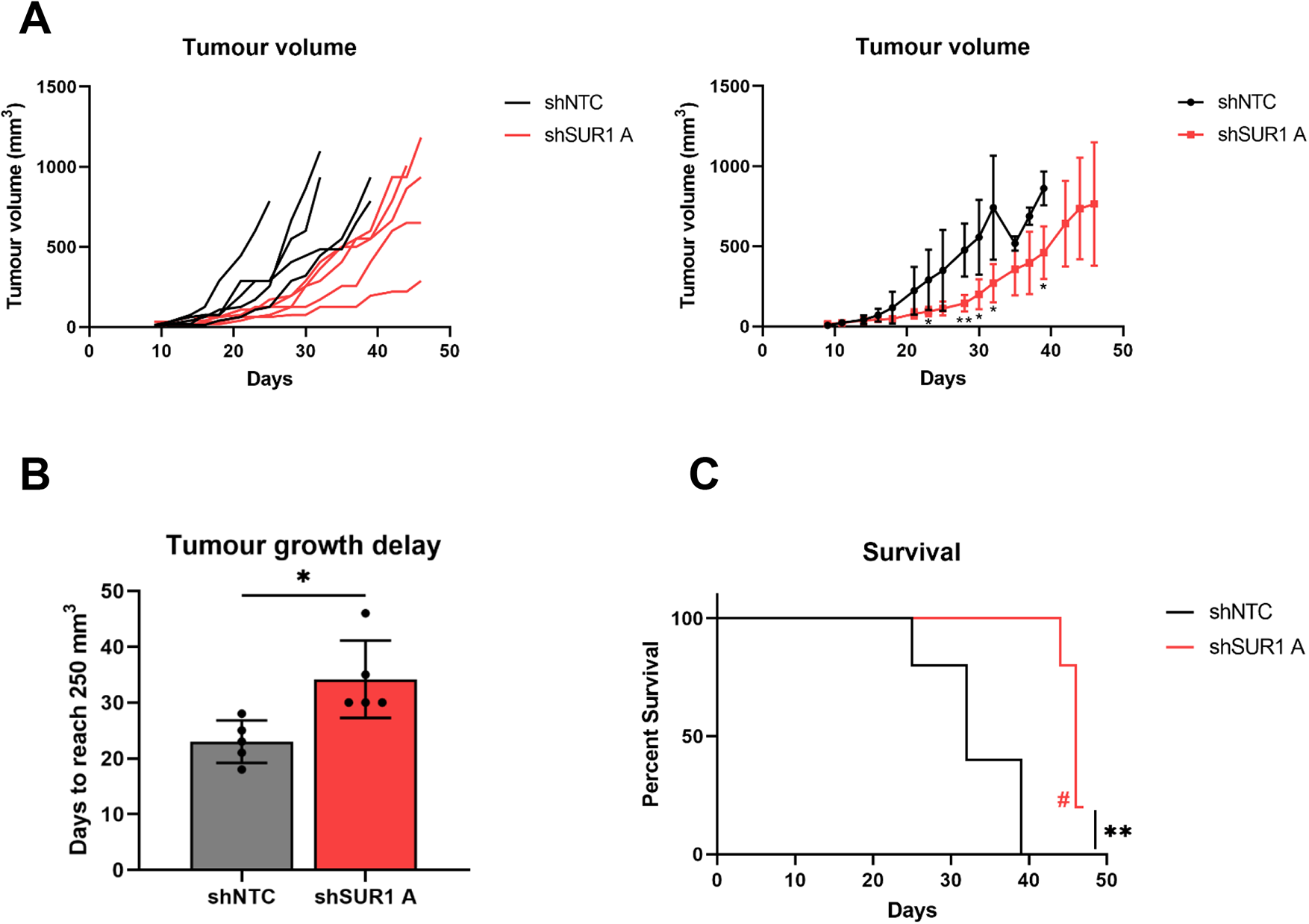
K_ATP_ channels drive proliferation in an *in vivo* mouse model. **A)** Tumour growth curves for mice implanted with HeLa cells stably expressing either a non-targetting (shNTC) or an SUR1-specific shRNA (shSUR1 A). Tumour volume was calculated using the formula V=0.5*L*W2. Both individual curves for each replicate (left) and curves representing mean values ± SD of five mice per group (right) are displayed. **B)** Tumour growth delay, calculated as the period of time between injection of tumours and growth to a set volume (250 mm^3^). Bars represent means ± SD of five biological replicates with individual data points displayed. *P<0.05 (Student’s t-test). **C)** Kaplan-Meier survival curve of mice bearing shNTC and shSUR1 A tumours. # indicates that one mice remained alive at the conclusion of the study. **P<0.01 (log-rank (Mantel-Cox) test).

## Discussion

It is vital to identify virus-host interactions that are critical for HPV-mediated transformation as, despite the availability of prophylactic vaccines, there are currently no effective anti-viral treatments for HPV-associated disease. Here, we identify a novel host factor, the ATP-sensitive potassium ion (K_ATP_) channel, as a crucial driver of cell proliferation in HPV+ cervical cancer (**Fig 10**). Inhibition of K_ATP_ channels, through either siRNA-mediated knockdown of individual subunits or pharmacological blockade using licenced inhibitors, significantly impedes proliferation and cell cycle progression. HPV is able to promote K_ATP_ channel activity via E7-mediated upregulation of the SUR1 subunit; this is observed in both cervical disease and cervical cancer tissue, as well as *in vitro* primary cell culture models of the HPV life cycle. As such, we believe that the clinically available inhibitors of K_ATP_ channels could constitute a potential novel therapy for HPV-associated malignancies.

**Fig 10.**
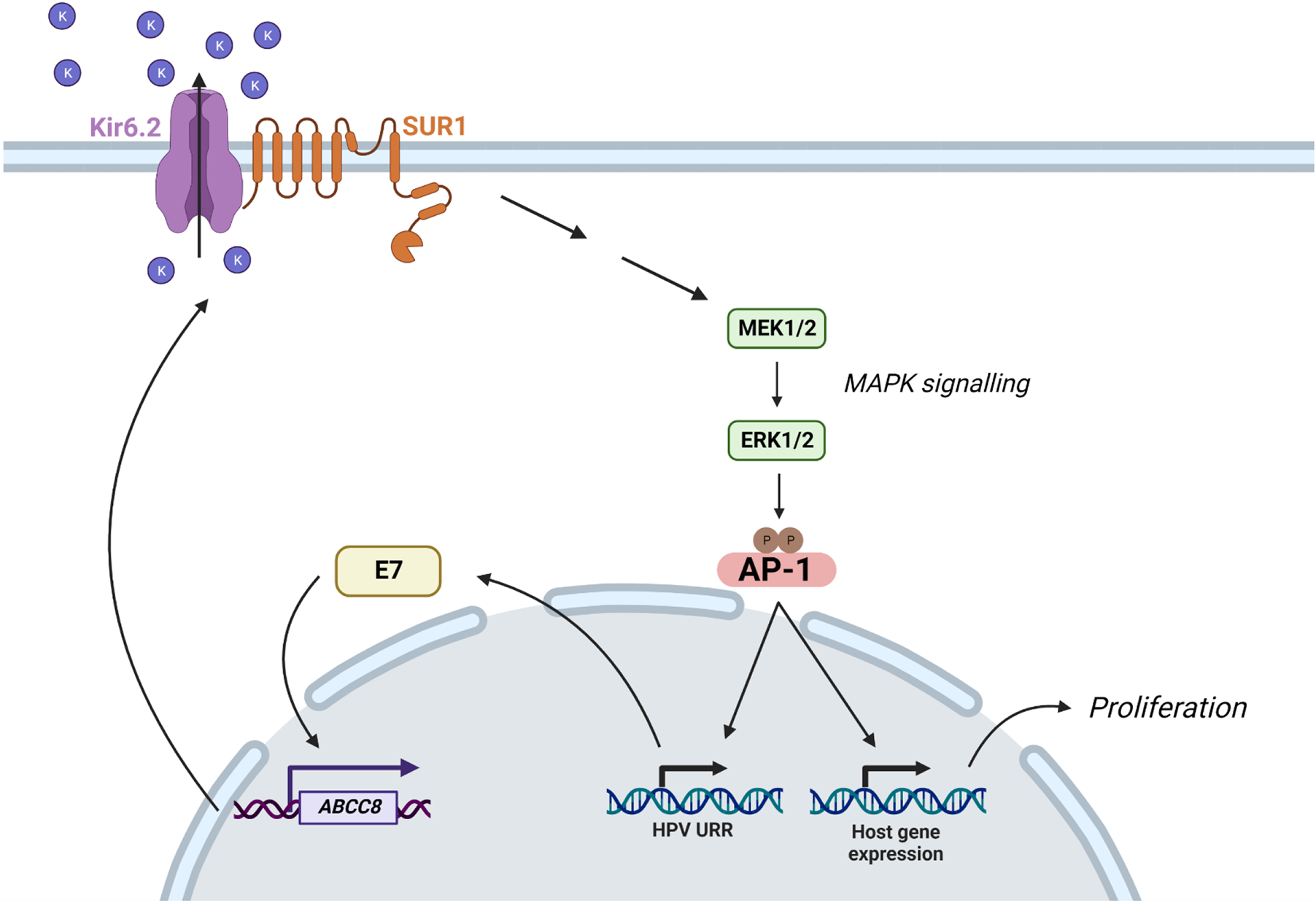
Schematic demonstrating E7-mediated upregulation of K_ATP_ channel expression and activity. HPV E7 upregulates expression of *ABCC8*, the gene encoding SUR1 which constitutes the regulatory subunit of K_ATP_ channels. Increased K_ATP_ channel activity contributes towards the activation of MAPK and AP-1 signalling. This, in turn, drives transcription from the viral URR and changes in host gene expression which together stimulate proliferation. Figure created using BioRENDER.com.

A growing number of viruses have been shown to modulate or require the activity of host ion channels [38]. Indeed, several viruses encode their own ion channels, termed ‘viroporins’, including the HPV E5 protein [57–59]. Together, this underlines the importance of regulating host ion channel homeostasis during infection. Until recently however, much research in this field had focussed on RNA viruses, but recent work in our laboratories have highlighted roles for host chloride, potassium and calcium ion channels during BK polyomavirus (BKPyV), Merkel cell polyomavirus (MCPyV) and Kaposi’s sarcoma-associated herpesvirus (KSHV) infection [60–62]. Importantly, this study is the first to our knowledge to explicitly demonstrate modulation of ion channel activity by HPV, and that this can contribute to host cell transformation. Although a previous study indicated that K_ATP_ channels are expressed in cervical cancer, no attempt was made to attribute this to HPV-mediated upregulation and the K_ATP_ channels found to be present were in fact comprised of Kir6.2 and the alternative regulatory subunit SUR2 [33]. Here, we found no evidence for expression of the SUR2A isoform, and SUR2B expression was not significantly increased in any of the HPV+ cell lines examined. Further, our functional analyses demonstrated that SUR2B has no impact on the proliferation of HPV+ cervical cancer cells.

More widely, few reports exist of a dependence on host K_ATP_ channel activity for viral replication. One study identified that inhibition of K_ATP_ channels via glibenclamide treatment precludes HIV cell entry but, in contrast, cardiac K_ATP_ channel activity was found to be detrimental to Flock House virus (FHV) infection of *Drosophila* [63, 64]. Significantly however, no evidence exists to suggest that either of these viruses actively modulate the gating and/or expression of these channels, as we have demonstrated for HPV here.

In order to confirm that the effects on HPV oncoprotein expression and proliferation observed during this study following SUR1 knockdown were due to decreased K_ATP_ channel activity, silencing of the pore-forming Kir6.2 subunit was also performed. We felt this pertinent as a recent studies concluded that the oncogenic activities of SUR1 in non-small cell lung carcinoma (NSCLC) are independent of K_ATP_ channel activity [65, 66], and SUR1 is reported to have a supplementary role in regulating the activity of an ATP-sensitive, non-selective ion channel in astrocytes [67]. However, we observed almost identical effects on cell proliferation and HPV oncoprotein expression following Kir6.2 knockdown, leading us to conclude that SUR1 does not act in a K_ATP_ channel-independent manner in cervical cancer. This is supported by our overexpression data, in which a significant increase in the proliferative ability of HPV-C33A cells was only observed when both channel subunits were transfected. This also fits with the current assembly hypothesis for K_ATP_ channels, whereby neither subunit can be trafficked beyond the endoplasmic reticulum unless fully assembled into hetero-octameric channels [68–70].

We observed that inhibition of K_ATP_ channels, through either pharmacological means or SUR1 knockdown, resulted in an increase in the proportion of cells in G1 phase and, consistently, a decrease in cyclin D1 and E1 expression. This is in line with prior reports in other cell types [30, 31, 49]. Further, this fits with an increasing recognition of the importance of ion channels in the regulation of the cell cycle and cell proliferation [24–26, 28]. It is thought that cells undergo a rapid hyperpolarisation during progression through the G1-S phase checkpoint, for which K^+^ efflux channels are particularly important [25, 26]. Our data indicates that, at least in cervical cancer, K_ATP_ channels may contribute towards this hyperpolarisation event. Further, several K^+^ channels demonstrate cell cycle-dependent variations in expression and/or activity [71–74]; whether this is the case for K_ATP_ channels in HPV+ cancer cells warrants further analysis.

Ion channels represent ideal candidates for novel cancer therapeutics given the abundance of licensed and clinically available drugs targeting the complexes which could be repurposed if demonstrated to be effective [23]. We therefore investigated whether K_ATP_ channel inhibition had a cytotoxic effect on cervical cancer cells. Somewhat surprisingly, given the impact on HPV oncoprotein expression, we did not observe any evidence for increased cell death following glibenclamide treatment. This is in contrast to previous experiments in gastric cancer, glioma and hepatocellular carcinoma cell lines [29, 31, 32], yet in agreement with observations in breast cancer cells [30]. These differences may potentially reflect the cell type-specific roles of K_ATP_ channels, or perhaps be a result of differing subunit compositions in the cell types analysed. Following this, we performed *in vivo* tumourigenicity assays using cell lines stably expressing SUR1-specific shRNAs. We observed significant delays to tumour growth with cells displaying reduced SUR1 expression resulting in prolonged survival, thus providing validation for our earlier *in vitro* work. Given the clear impact suppression of K_ATP_ channel activity has on the growth of HPV+ cervical cancer cells, further work is now warranted to confirm whether the licenced K_ATP_ channel inhibitors could be repurposed and used alongside current therapies.

We revealed that the pro-proliferative effects of K_ATP_ channels are mediated via activation of ERK1/2 and subsequently the AP-1 family member cJun. Previous reports have examined the importance of these signalling pathways in HPV infection and cervical cancer [53, 75–79]. Indeed, a recent study elegantly showed a strong correlation between ERK1/2 activity and cervical disease progression, and highlighted the importance of ERK1/2 and AP-1 signalling for oncoprotein expression in both a life cycle model of HPV infection and using an oropharyngeal squamous cell carcinoma cell line [79]. Interestingly, this study additionally identified the AP-1 family members cFos and JunB as contributors towards oncogene transcription, whilst our own analysis has revealed that both cJun and JunD are upregulated in HPV18+ keratinocytes and cervical cancer cell lines [53, 79]. As AP-1 can be comprised of Jun family homodimers or heterodimers with Fos, ATF or MAF family proteins, further studies may be warranted to determine the most frequent makeup of AP-1 dimers in HPV+ cells [51].

In conclusion, we present evidence that host K_ATP_ channels play a crucial role in cervical carcinogenesis. Upregulation of the SUR1 subunit by HPV E7 contributes towards increased K_ATP_ channel activity, which in turn drives cell proliferation and progression through the G1/S phase checkpoint via MAPK/AP-1 signalling. K_ATP_ channels also promote HPV E6/E7 expression, thus establishing a positive feedback network. A complete characterisation of the role of K_ATP_ channels in HPV-associated disease is therefore now warranted in order to determine whether the licensed and clinically available inhibitors of these channels could constitute a potential novel therapy in the treatment of HPV-driven cancers.

## Materials and Methods

### Cervical cancer cytology samples

Cervical cytology samples were obtained from the Scottish HPV Archive (http://www.shine.mvm.ed.ac.uk/archive.shtml), a biobank of over 20,000 samples designed to facilitate HPV-associated research. The East of Scotland Research Ethics Service has given generic approval to the Scottish HPV Archive as a Research Tissue Bank (REC Ref 11/AL/0174) for HPV related research on anonymised archive samples. Samples are available for the present project though application to the Archive Steering Committee (HPV Archive Application Ref 0034). RNA was extracted from the samples using TRIzol® Reagent (ThermoFisher Scientific) and analysed as described.

### Plasmids and siRNA

Expression vectors for ΔJunD, HA-tagged Kir6.2 and SUR1, and GFP-tagged HPV18 E6 and E7 have been described previously [53, 80, 81]. The FLAG-tagged HPV18 E7 expression vector was cloned from the above GFP-HPV18 E7 vector. Luciferase reporter constructs for the HPV18 URR and the HPV16 URR were kind gifts from Prof Felix Hoppe-Seyler (German Cancer Research Center (DKFZ)) and Dr Iain Morgan (Virginia Commonwealth University (VCU)) respectively [76, 82]. The HA-tagged cJun S63/73D expression vector was kindly provided by Dr Hans van Dam (Leiden University Medical Centre (LUMC)) [83]. The AP-1 luciferase reporter has been described previously [52].

For siRNA experiments, pools of four siRNAs specific to *ABCC8* (FlexiTube GeneSolution GS6833), *ABCC9* (FlexiTube GeneSolution GS10060), and *KCNJ11* (FlexiTube GeneSolution GS3767) were purchased from Qiagen. HPV16 E6 siRNA (sc-156008) and HPV16 E7 siRNA (sc-270423) were purchased from Santa Cruz Biotechnology (SCBT). HPV18 E6 siRNAs were purchased from Dharmacon (GE Healthcare) and had the following sequences: 5’-CUAACACUGGGUUAUACAA-3’ and 5’-CTAACTAACACTGGGTTAT-3’. HPV18 E7 siRNA were as previously described and were a kind gift from Prof Eric Blair (University of Leeds) [84, 85]. A final siRNA concentration of 25 nM was used in all cases.

### K^+^ channel modulators and small molecule inhibitors

The K_ATP_ inhibitors glibenclamide and tolbutamide were purchased from Sigma and used at final concentrations of 10 μM and 200 μM unless stated otherwise. The K^+^ channel blockers tetraethylammonium (TEA), quinine, quinidine, 4-aminopyridine (4-AP), ruthenium red (RR), apamin, bupivacaine hydrochloride (BupHCl) and margatoxin were purchased from Sigma and used at the stated concentrations. The K_ATP_ channel activator diazoxide was purchased from Cayman Chemical and used at 50 μM unless stated otherwise. The MEK1/2 inhibitor U0126 (Calbiochem) was used at 20 μM. Staurosporine (Calbiochem) was used at a final concentration of 1 μM.

### Cell culture

HeLa (HPV18+ cervical epithelial adenocarcinoma cells), SW756 (HPV18+ cervical squamous carcinoma cells), SiHa (HPV16+ cervical squamous carcinoma cells), CaSki (HPV16+ cervical squamous carcinoma cells) and C33A (HPV-cervical squamous carcinoma) cells obtained from the ATCC were grown in DMEM supplemented with 10% FBS (ThermoFisher Scientific) and 50 U/mL penicillin/streptomycin (Lonza). HEK293TT cells were kindly provided by Prof Greg Towers (University College London (UCL)) and grown as above.

Neonate foreskin tissues were obtained from local General Practice surgeries and foreskin keratinocytes isolated under ethical approval no 06/Q1702/45. Cells were maintained in serum-free medium (SFM; Gibco) supplemented with 25 µg/mL bovine pituitary extract (Gibco) and 0.2 ng/mL recombinant EGF (Gibco). The transfection of primary NHKs to generate HPV18+ keratinocytes was performed as described previously [86]. All cells were cultured at 37 °C and 5% CO_2_.

All cells were negative for mycoplasma during this investigation. Cell identity was confirmed by STR profiling.

### Organotypic raft culture

NHKs and HPV18+ keratinocytes were grown in organotypic raft cultures by seeding the keratinocytes onto collagen beds containing J2-3T3 fibroblasts [86]. Once confluent, the collagen beds were transferred onto metal grids and fed from below with FCS-containing E media without EGF. The cells were allowed to stratify for 14 days before fixing with 4% formaldehyde. The rafts were paraffin-embedded and 4 μm tissue sections prepared (Propath UK Ltd.). For analysis of SUR1 expression, the formaldehyde-fixed raft sections were treated with the sodium citrate method of antigen retrieval. Briefly, sections were boiled in 10 mM sodium citrate with 0.05% Tween-20 for 10 minutes. Sections were incubated with a polyclonal antibody against SUR1 (PA5-50836, ThermoFisher Scientific) and immune complexes visualised using Alexa 488 and 594 secondary antibodies (Invitrogen). The nuclei were counterstained with DAPI and mounted in Prolong Gold (Invitrogen).

### Transfection of cancer cell lines

Transient transfections were performed using Lipofectamine 2000 (ThermoFisher Scientific). A ratio of nucleic acid to Lipofectamine 2000 of 1:2 was used for both DNA and siRNA. Transfections were performed overnight in OptiMEM I Reduced Serum Media (ThermoFisher Scientific).

### Generation of stable cell lines

HEK293TT cells were co-transfected with the packaging plasmids pCRV1-NLGP and pCMV-VSV-G alongside either a pZIP-hEF1α-non-targeting shRNA construct or one of three pZIP-hEF1α-SUR1 shRNA constructs (purchased from TransOMIC). At 48 hours post-transfection, virus-containing media was harvested and passed through a 0.45 μm filter to remove cell debris. To perform lentiviral transduction, culture media was removed from HeLa cells seeded 24 hours earlier and replaced with virus-containing media. Cells were incubated overnight before removing virus and replacing with complete DMEM. At 48 hours post-transduction, cells were passaged as appropriate and treated with 1 μg/mL puromycin in culture media for 48 hours to select for transduced cells. Fluorescence-associated cell sorting (FACS) was used to partition individual surviving cells into wells of 96-well culture plates in order to generate monoclonal cell lines.

HPV-C33A cells stably expressing HPV18 E6 or E7 were generated as previously described [18].

### Western blot analysis

Equal amounts of protein from cell lysates were resolved by molecular weight using 8-15% SDS-polyacrylamide gels as appropriate. Separated proteins were transferred to Hybond™ nitrocellulose membranes (GE Healthcare) using a semi-dry method (Bio-Rad Trans-Blot® Turbo™ Transfer System). Membranes were blocked in 5% skimmed milk powder in tris-buffered saline-0.1% Tween 20 (TBS-T) for 1 hour at room temperature before probing with antibodies specific for HPV16 E6 (GTX132686, GeneTex, Inc.), HPV16 E7 (ED17: sc-6981, SCBT), HPV18 E6 (G-7: sc-365089, SCBT), HPV18 E7 (8E2: ab100953, abcam), HA (3724, Cell Signalling Technology (CST)), cyclin A (B-8: sc-271682, SCBT), cyclin B1 (12231, CST), cyclin D1 (ab134175, abcam), cyclin E1 (20808, CST), PARP (9542, CST), caspase-3 (9662, CST), cleaved caspase-3 (D175) (9664, CST), phospho-ERK1/2 (T202/Y204) (9101, CST), ERK1/2 (9102, CST), phospho-cJun (S73) (3270, CST), cJun (9165, CST), JunD (5000, CST) and GAPDH (G-9: sc-365062, SCBT). Primary antibody incubations were performed overnight at 4 °C. The appropriate HRP-conjugated secondary antibodies (Jackson ImmunoResearch) were used at a 1:5000 dilution. Blots were visualised using ECL reagents and CL-XPosure™ film (ThermoFisher Scientific).

### RNA extraction and reverse transcription-quantitative PCR (RT-qPCR)

Total RNA was extracted from cells using the E.Z.N.A.® Total RNA Kit I (Omega Bio-Tek) following the provided protocol for RNA extraction from cultured cells. The concentration of eluted RNA was determined using a NanoDrop™ One spectrophotometer (ThermoFisher Scientific). RT-qPCR was performed using the GoTaq® 1-Step RT-qPCR System (Promega) with an input of 50 ng RNA. Reactions were performed using a CFX Connect Real-Time PCR Detection System (BioRad) with the following cycling conditions: reverse transcription for 10 min at 50 °C; reverse transcriptase inactivation/polymerase activation for 5 min at 95 °C followed by 40 cycles of denaturation (95 °C for 10 sec) and combined annealing and extension (60 °C for 30 sec). Data was analysed using the ΔΔCt method [87]. The primers used in this study are detailed in S1 Table; *U6* expression was used for normalisation.

### Tissue microarray and immunohistochemistry

A cervical cancer tissue microarray (TMA) containing 39 cases of cervical cancer and 9 cases of normal cervical tissue (in duplicate) was purchased from GeneTex, Inc. (GTX21468). Slides were deparaffinised in xylene, rehydrated in a graded series of ethanol solutions and subjected to antigen retrieval in citric acid. Slides were blocked in normal serum and incubated in primary antibody (SUR1 (75-267, Antibodies Inc.)) overnight at 4 °C. Slides were then processed using the VECTASTAIN^®^ Universal Quick HRP Kit (PK-7800; Vector Laboratories) as per the manufacturer’s instructions. Immunostaining was visualised using 3,3’-diaminobenzidine (Vector^®^ DAB (SK-4100; Vector Laboratories)). Images were taken using an EVOS® FL Auto Imaging System (ThermoFisher Scientific) at 20x magnification. SUR1 quantification was automated using ImageJ with the IHC Profiler plug-in [88, 89]. Histology scores (H-score) were calculated based on the percentage of positively stained tumour cells and the staining intensity grade. The staining intensities were classified into the following four categories: 0, no staining; 1, low positive staining; 2, positive staining; 3, strong positive staining. H-score was calculated by the following formula: (3 x percentage of strong positive tissue) + (2 x percentage of positive tissue) + (percentage of low positive tissue), giving a range of 0 to 300.

### Patch clamping

HeLa cells were seeded on coverslips in 12-well culture plates at 10-20% confluency to prevent cell-cell contact. Following attachment, cells were treated with DMSO, 10 μM glibenclamide, 50 μM diazoxide, or with both channel modulators in combination for 16 hours. Following treatment, patch pipettes (2–4 MΩ) were filled with pipette solution (5 mM HEPES-KOH pH 7.2, 140 mM KCl, 1.2 mM MgCl_2_, 1 mM CaCl_2_, 10 mM EGTA, 1 mM MgATP, 0.5 mM NaUDP) and culture media removed from cells and replaced with external solution (5mM HEPES-KOH pH 7.4, 140 mM KCl, 2.6 mM CaCl_2_, 1.2 mM MgCl_2_). Whole cell patch clamp recordings were performed using an Axopatch 200B amplifier/Digidata 1200 interface controlled by Clampex 9.0 software (Molecular Devices). A series of depolarising steps, from −100 to +60 mV in 10 mV increments for 100 ms each, was applied to cells and the K^+^ current measured. Analysis was performed using the data analysis package Clampfit 9.0 (Molecular Devices).

### Chromatin immunoprecipitation (ChIP)

After treatment as required, cells were processed for ChIP as previously described [81]. cJun was immunoprecipitated using a ChIP grade anti-cJun antibody (9165, CST) and each of the samples were also pulled down with an IgG isotype control to confirm antibody specificity. Pierce™ Protein A/G Magnetic Beads (ThermoFisher Scientific) were used to isolate antibody-chromatin complexes. Immunoprecipitated chromatin was then processed for quantitative PCR (qPCR) using primers covering the AP-1 binding sites within the HPV18 URR (sequences available upon request). Fold enrichment was calculated by comparing to the IgG isotype control.

### Luciferase reporter assays

Cells were transfected with plasmids expressing the appropriate firefly luciferase reporter (250 ng). 25 ng of a *Renilla* luciferase reporter construct (pRLTK) was used as an internal control for transfection efficiency. Samples were lysed in passive lysis buffer (Promega) and activity measured using a dual-luciferase reporter assay system (Promega). All assays were performed in triplicate, and each experiment was repeated a minimum of three times.

### Proliferation assays

For cell growth assays, cells were detached by trypsinisation after treatment as necessary and reseeded at equal densities in 12-well plates. Cells were subsequently harvested every 24 hours and manually counted using a haemocytometer.

For colony formation assays, cells were detached by trypsinisation after treatment as required and reseeded at 500 cells/well in six-well plates. Once visible colonies were noted, culture media was aspirated and cells fixed and stained in crystal violet staining solution (1% crystal violet, 25% methanol) for 15 min at room temperature. Plates were washed thoroughly with water to remove excess crystal violet and colonies counted manually.

For soft agar assays, 60 mm cell culture plates were coated with a layer of complete DMEM containing 0.5% agarose. Simultaneously, cells were detached by trypsinisation after treatment as required and resuspended at 1000 cells/mL in complete DMEM containing 0.35% agarose and added to the bottom layer of agarose. Once set, plates were covered with culture media and incubated for 14-21 days until visible colonies were observed. Colonies were counted manually.

### Cell cycle analysis

Cells were harvested and fixed overnight in 70% ethanol at −20°C. Ethanol was removed by centrifugation at 500 x g for 5 min and cells washed twice in PBS containing 0.5% BSA. Cells were resuspended in 500 μL 0.5% BSA/PBS, treated with 1.25 μL RNase A/T1 mix (ThermoFisher Scientific) and stained with 8 μL 1 mg/mL propidium iodide solution (Sigma) for 30 min at room temperature in the dark. Analysis was performed using a CytoFLEX S flow cytometer (Beckman Coutler).

### DiBAC assay

After treatment as necessary, the membrane potential-sensitive dye DiBAC_4_(3) (Bis-(1,3-Dibutylbarbituric Acid) Trimethine Oxonol; ThermoFisher Scientific) was added directly to culture media at a final concentration of 200 nM. Cells were incubated in the presence of the dye for 20 min at 37°C in the dark. Cells were harvested by scraping and washed twice in PBS. Cells were resuspended in 500 μL PBS for flow cytometry analysis. Analysis was performed using a CytoFLEX S flow cytometer (Beckman Coutler).

### Annexin V assay

Annexin V apoptosis assays were performed using the TACS® Annexin V-FITC Kit (Bio-Techne Ltd.). After treatment as required, cells were harvested by aspirating and retaining culture media (to collect detached apoptotic cells) with the remaining cells detached by trypsinisation. The retained media and trypsin cell suspension was combined and centrifuged at 500 x g for 5 min to pellet cells before washing once in PBS and pelleting again. Cells were incubated in 100 μL Annexin V reagent (10 μL 10X binding buffer, 10 μL propidium iodide, 1 μL Annexin V-FITC (diluted 1:25), 79 μL ddH_2_O) for 15 min at room temperature protected from light. 400 μL of 1X binding buffer was then added before analysing using a CytoFLEX S flow cytometer (Beckman Coutler). Annexin V-FITC positive cells were designated as early apoptotic, whilst dual Annexin V-FITC/PI positive cells were designated as late apoptotic. Cells negative for both Annexin V and PI staining were considered to be healthy.

### *In vivo* tumourigenicity study

Female 6-8 week old SCID mice were purchased from Charles River Laboratories. All animal work was carried out under project license PP1816772. HeLa cells stably expressing either a non-targeting shRNA or a SUR1-specific shRNA were harvested, pelleted and resuspended in sterile PBS. Five mice were used per experimental group, with each injected subcutaneously with 5 x 10^5^ cells in 50 µL PBS. Once palpable tumours had formed (~10 days), measurements for both groups were taken thrice weekly. After tumours reached 10 mm in either dimension, mice were monitored daily. Mice were sacrificed once tumours reached 15 mm in any dimension. No toxicity, including significant weight loss, was seen in any of the mice. Tumour volume was calculated with the formula V = 0.5*L*W^2^.

### Microarray analysis

For microarray analysis, a dataset consisting of 24 normal, 14 CIN1 lesions, 22 CIN2 lesions, 40 CIN3 lesions, and 28 cancer specimens was utilised. Microarray data was obtained from GEO database accession number GSE63514 [90].

### Statistical analysis

All experiments were performed a minimum of three times, unless stated otherwise. Data was analysed using a two-tailed, unpaired Student’s t-test performed using GraphPad PRISM 9.2.0 software, unless stated otherwise. Kaplan-Meier survival data was analysed using the log-rank (Mantel-Cox) test.

## Acknowledgements

We are grateful to Prof Felix Hoppe-Seyler (DKFZ), Dr Iain Morgan (VCU), Prof Eric Blair (University of Leeds), Dr Hans van Dam (LUMC) and Prof Greg Towers (UCL) for provision of reagents. We thank the Scottish HPV Investigators Network (SHINE), Prof Sheila Graham (University of Glasgow), Dr David Millan (University of Glasgow) and Prof Nick Coleman (University of Cambridge) for providing HPV positive patient samples.

## Author Contributions

Conceptualisation (JAS, CWW, JM, ELM, AM); Formal analysis (JAS, CWW, MRP, HC, ELM); Funding acquisition (AW, AS, ELM, AM); Investigation (JAS, CWW, MRP, DE, HC, ELM); Project administration (AW, AS, AM); Resources (DBM); Supervision (AW, AS, AM); Writing – original draft (JAS); Writing – review & editing (all authors)

---

**S1 Fig.**
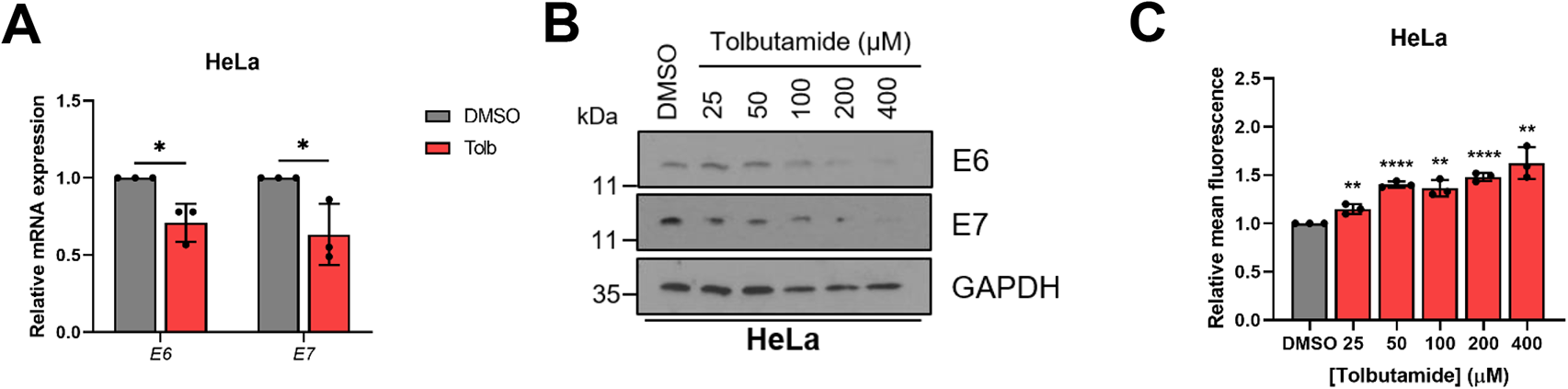
K_ATP_ channels are important for HPV gene expression in cervical cancer cells. **A)** Expression levels of *E6* and *E7* mRNA in HeLa cells treated with tolbutamide (200 μM) measured by RT-qPCR. Samples were normalised against *U6* mRNA levels. **B)** Representative western blots of E6 and E7 expression in HeLa cells treated with increasing doses of tolbutamide. GAPDH served as a loading control. **C)** Mean DiBAC_4_(3) fluorescence levels in HeLa cells treated with increasing dose of tolbutamide. Samples were normalised to DMSO control. Data represent means ± SD of a minimum of three biological replicates. *P<0.05, **P<0.01, ***P<0.001, ****P<0.0001 (Student’s t-test).

**S2 Fig.**
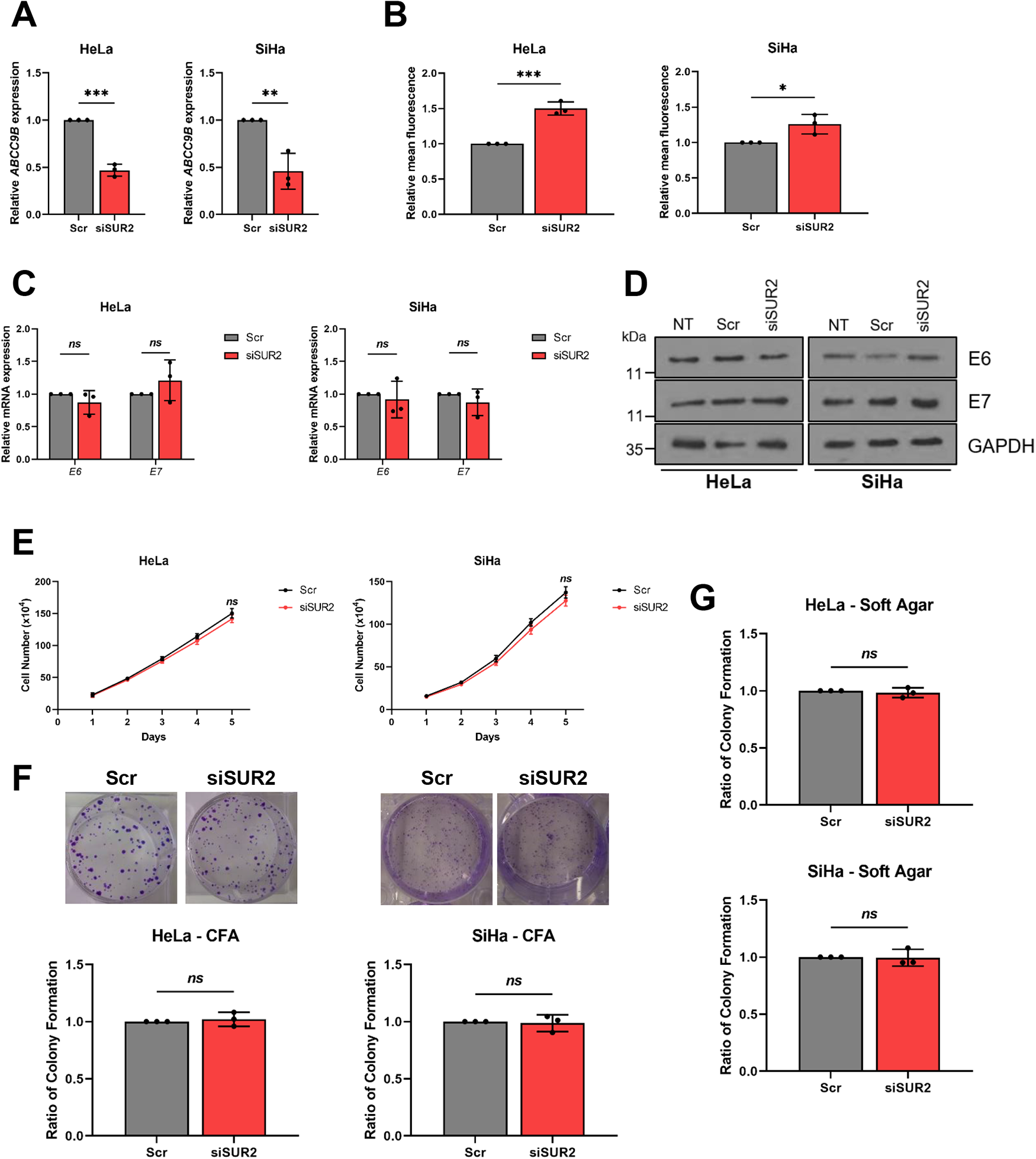
Depletion of SUR2 has no impact on HPV gene expression or proliferation in cervical cancer cells. **A)** Relative expression of *ABCC9B* mRNA in HeLa and SiHa cells transfected with a pool of SUR2-specific siRNA measured by RT-qPCR. Samples were normalised against *U6* mRNA levels. **B)** Relative mean DiBAC_4_(3) fluorescence levels in HeLa and SiHa cells transfected with SUR2 siRNA. **C)** Relative expression of *E6* and *E7* mRNA in HeLa and SiHa cells transfected with SUR2 siRNA measured by RT-qPCR. Samples were normalised against *U6* mRNA levels. **D)** Representative western blots of E6 and E7 expression in HeLa and SiHa cells transfected with SUR2 siRNA. GAPDH served as a loading control. **E-G)** Growth curve analysis **(E)**, colony formation assay **(F)** and soft agar assay **(G)** of HeLa and SiHa cells after transfection of SUR2-specific siRNA. Data represent means ± SD of a minimum of three biological replicates with individual data points displayed. *Ns* not significant, *P<0.05, **P<0.01, ***P<0.001 (Student’s t-test).

**S3 Fig.**
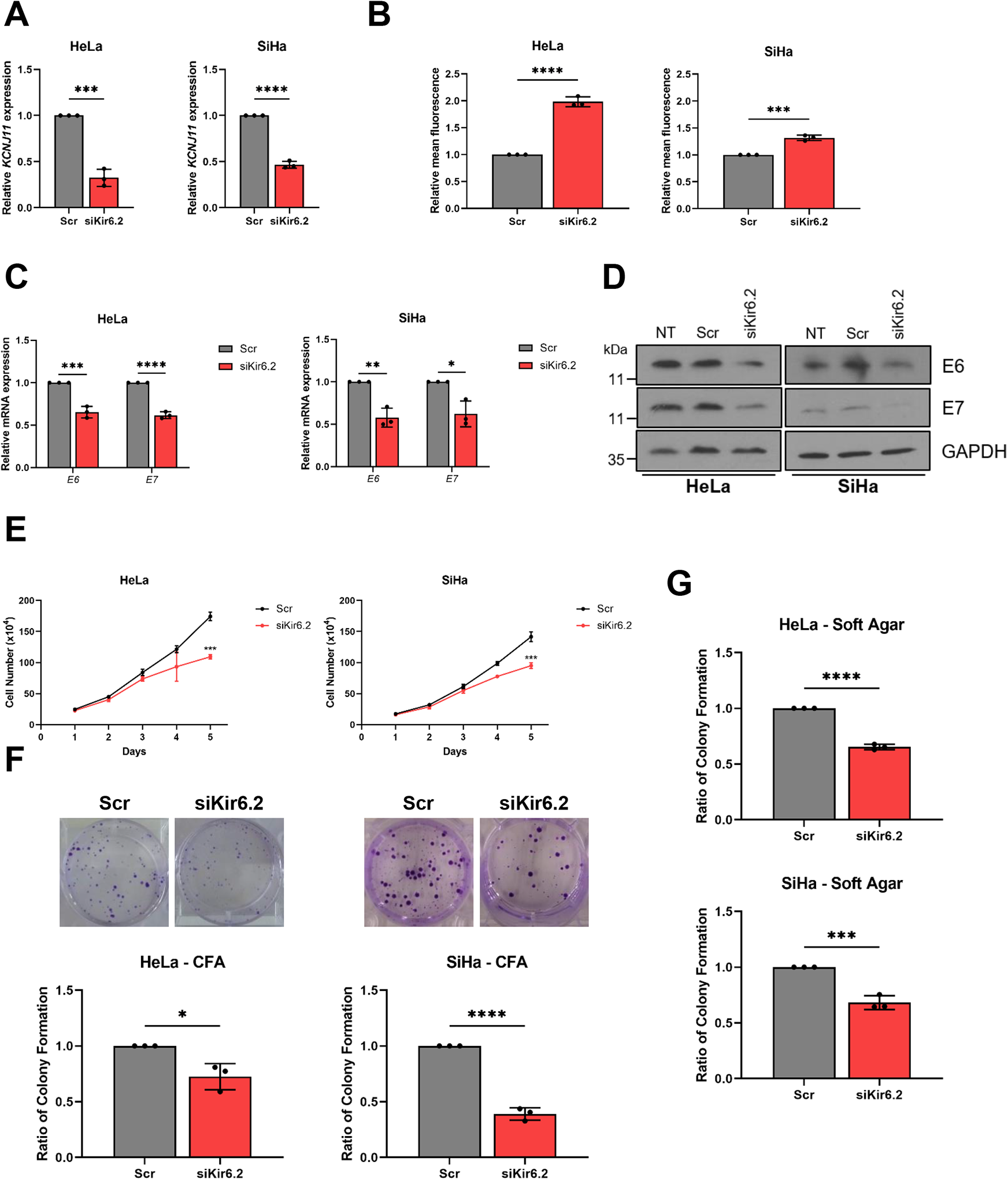
Depletion of Kir6.2 reduces HPV gene expression and proliferation in cervical cancer cells. **A)** Relative expression of *KCNJ11* mRNA in HeLa and SiHa cells transfected with a pool of Kir6.2-specific siRNA measured by RT-qPCR. Samples were normalised against *U6* mRNA levels. **B)** Relative mean DiBAC_4_(3) fluorescence levels in HeLa and SiHa cells transfected with Kir6.2 siRNA. **C)** Relative expression of *E6* and *E7* mRNA in HeLa and SiHa cells transfected with Kir6.2 siRNA measured by RT-qPCR. Samples were normalised against *U6* mRNA levels. **D)** Representative western blots of E6 and E7 expression in HeLa and SiHa cells transfected with Kir6.2 siRNA. GAPDH served as a loading control. **E-G)** Growth curve analysis **(E)**, colony formation assay **(F)** and soft agar assay **(G)** of HeLa and SiHa cells after transfection of Kir6.2-specific siRNA. Data shown is means ± SD of three biological replicates with individual data points displayed where appropriate. *P<0.05, **P<0.01, ***P<0.001, ****P<0.0001 (Student’s t-test).

**S4 Fig.**
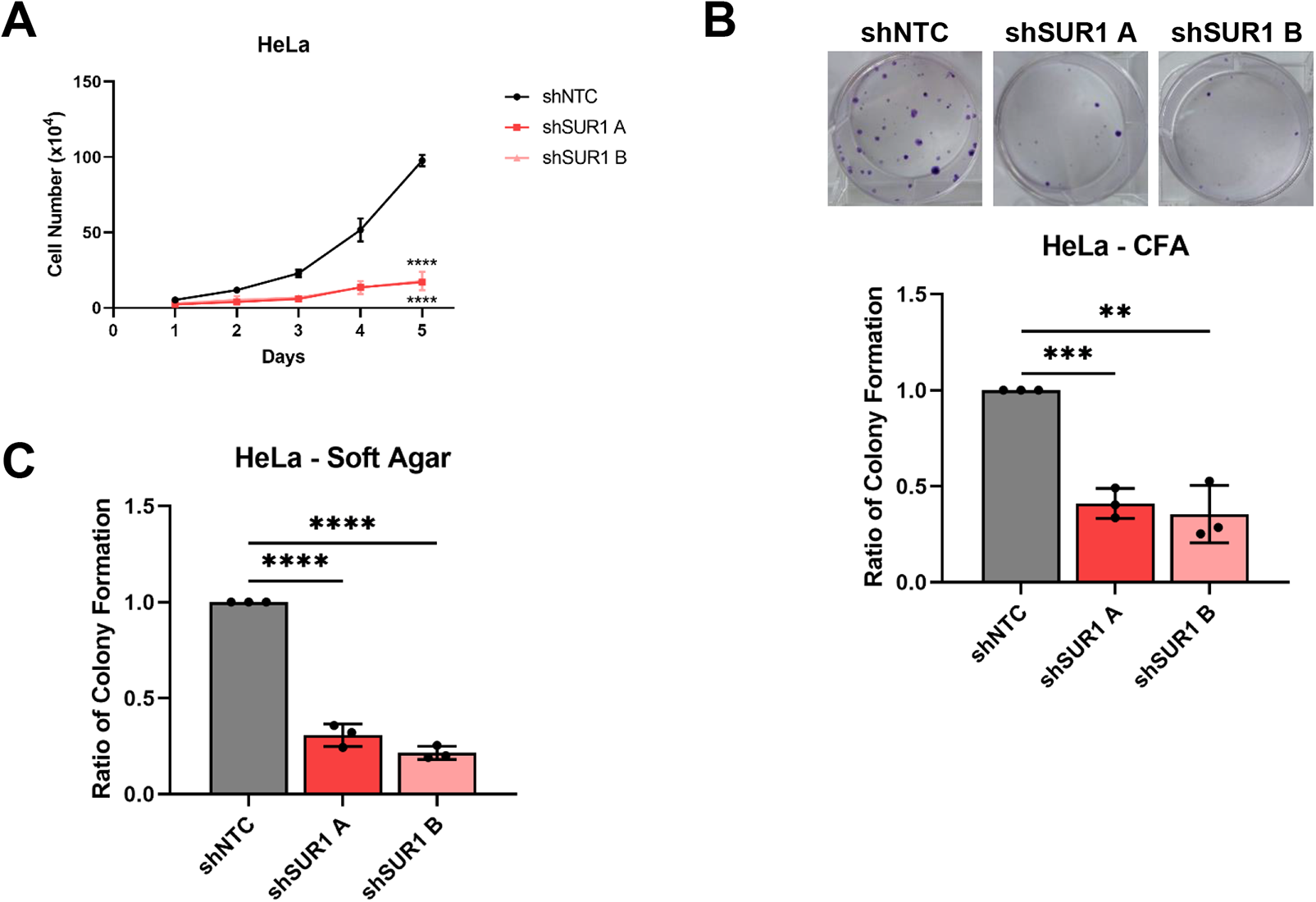
Stable suppression of SUR1 expression decreases the proliferation of cervical cancer cells. Growth curve analysis **(A)**, colony formation assay **(B)** and soft agar assay **(C)** of monoclonal HeLa cell lines stably expressing either a non-targeting (shNTC) or a SUR1-specific shRNA. Data shown is means ± SD of three biological replicates with individual data points displayed where appropriate. *P<0.05, **P<0.01, ***P<0.001, ****P<0.0001 (Student’s t-test).

**S5 Fig.**
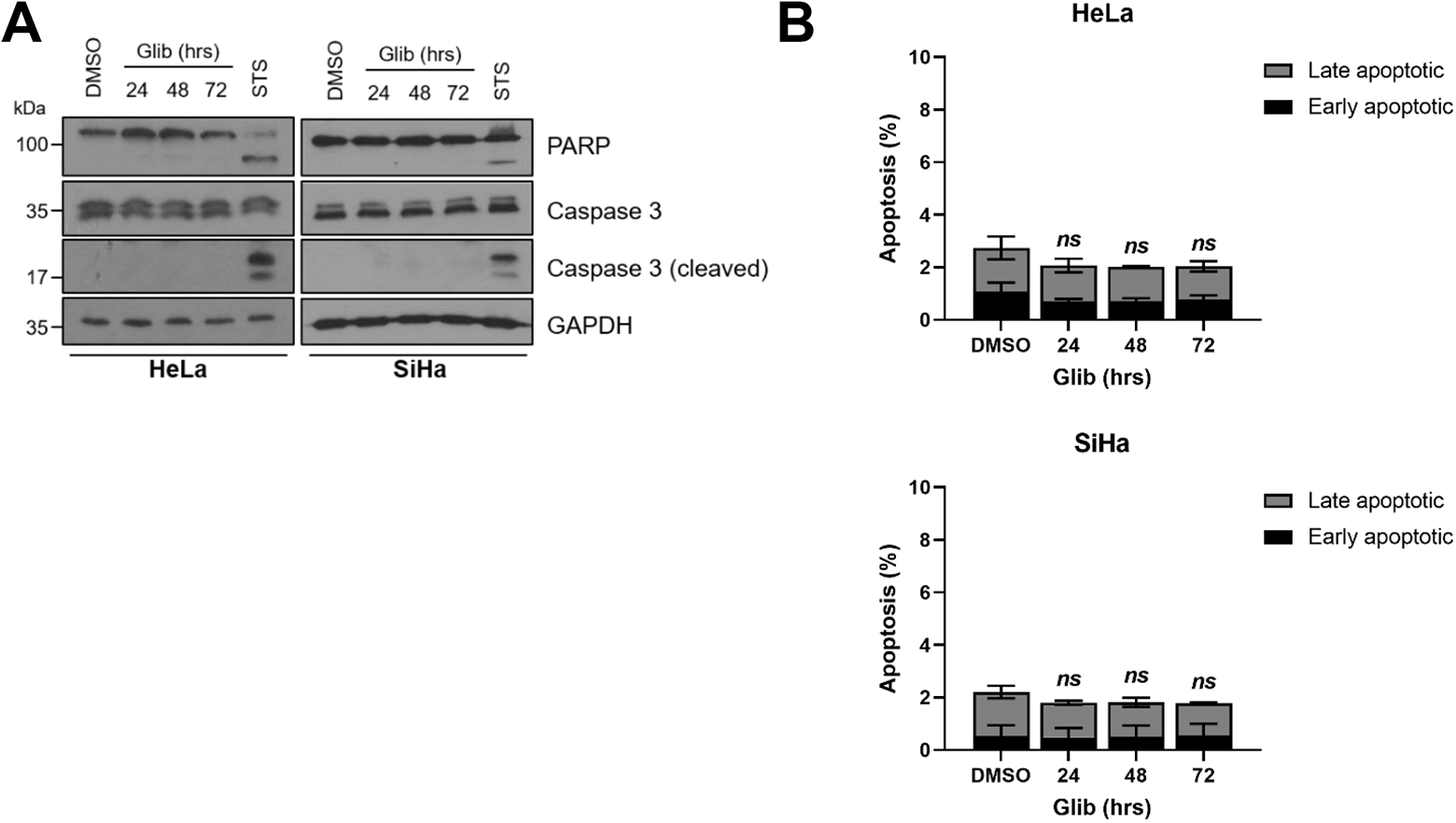
K_ATP_ channel inhibition does not impact upon the survival of cervical cancer cells. **A)** Representative western blots of PARP and caspase 3 cleavage in HeLa and SiHa cells treated with DMSO or glibenclamide (10 μM) for the indicated durations. Staurosporine treatment (STS, 1 μM for 6 hours) served as a positive control for apoptosis induction. GAPDH served as a loading control. **B)** Flow cytometry analysis of Annexin V assay using HeLa and SiHa cells treated with DMSO or glibenclamide (10 μM) for the indicated durations. Bars represent means ± SD of three biological replicates. *Ns* not significant (Student’s t-test).

**S1 Table.**
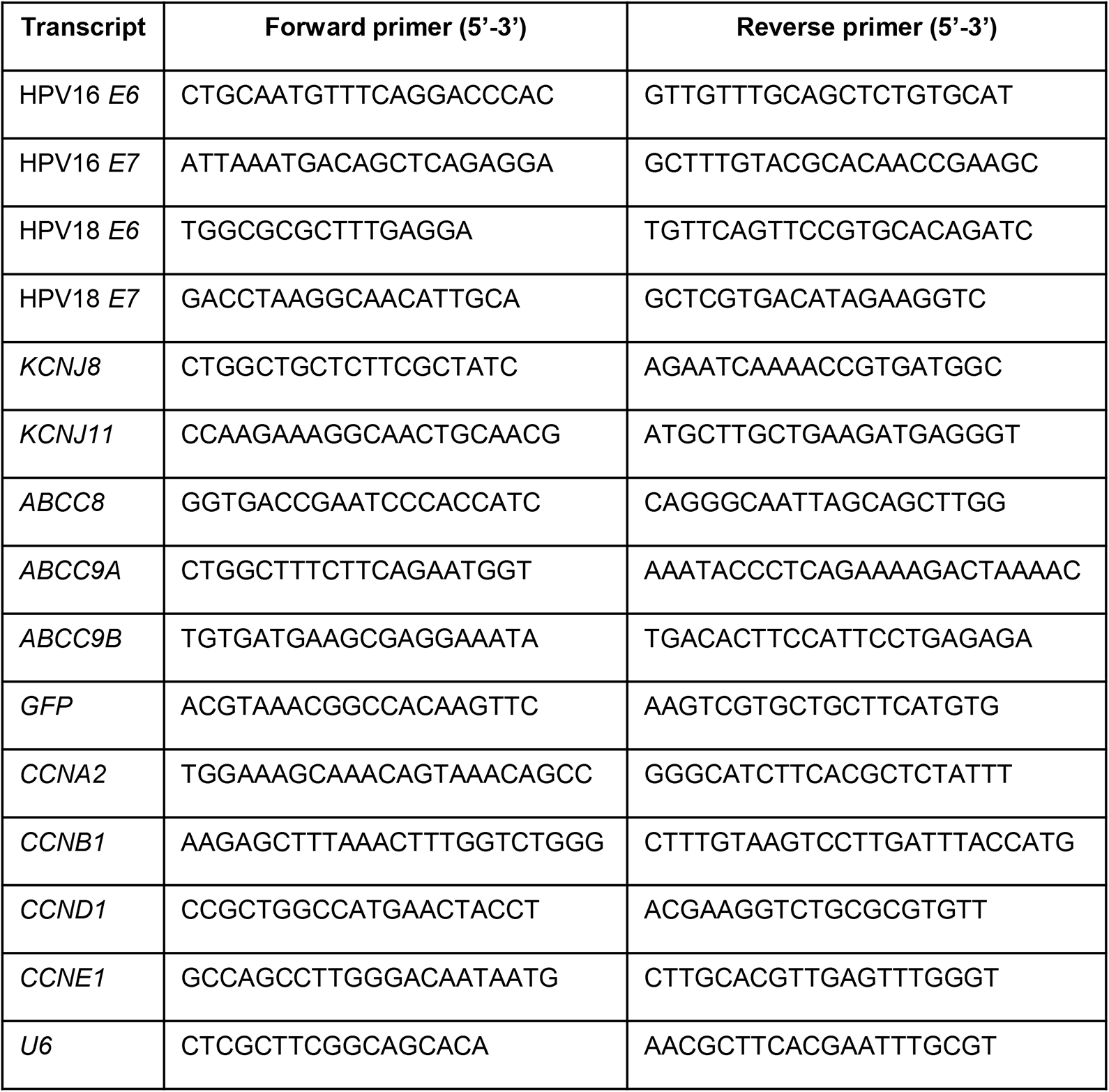
List of primers used for RT-qPCR in this study.

